# Translational control of polyamine metabolism by CNBP is required for Drosophila locomotor function

**DOI:** 10.1101/2021.04.29.441910

**Authors:** Sonia Coni, Federica A. Falconio, Marta Marzullo, Marzia Munafò, Benedetta Zuliani, Federica Mosti, Alessandro Fatica, Zaira Ianniello, Rosa Bordone, Alberto Macone, Enzo Agostinelli, Alessia Perna, Tanja Matkovic, Stephan Sigrist, Gabriella Silvestri, Gianluca Canettieri, Laura Ciapponi

## Abstract

Microsatellite expansions of CCTG repeats in the CNBP gene leads to accumulation of toxic RNA and have been associated to DM2. However, it is still unclear whether the dystrophic phenotype is also linked to CNBP decrease, a conserved CCHC-type zinc finger RNA binding protein that regulates translation and is required for mammalian development.

Here we show that depletion of *Drosophila* CNBP in muscles causes age-dependent locomotor defects that are correlated with impaired polyamine metabolism. We demonstrate that the levels of ornithine decarboxylase (ODC) and polyamines are significantly reduced upon dCNBP depletion. Of note, we show a reduction of the CNBP-polyamine axis in muscle from DM2 patients. Mechanistically, we provide evidence that dCNBP controls polyamine metabolism through binding dOdc mRNA and regulating its translation. Remarkably, the locomotor defect of dCNBP-deficient flies is rescued by either polyamine supplementation or dOdc1 overexpression. We suggest that this dCNBP function is evolutionarily conserved in vertebrates with relevant implications for CNBP-related pathophysiological conditions.

**GRAPHICAL ABSTRACT:** 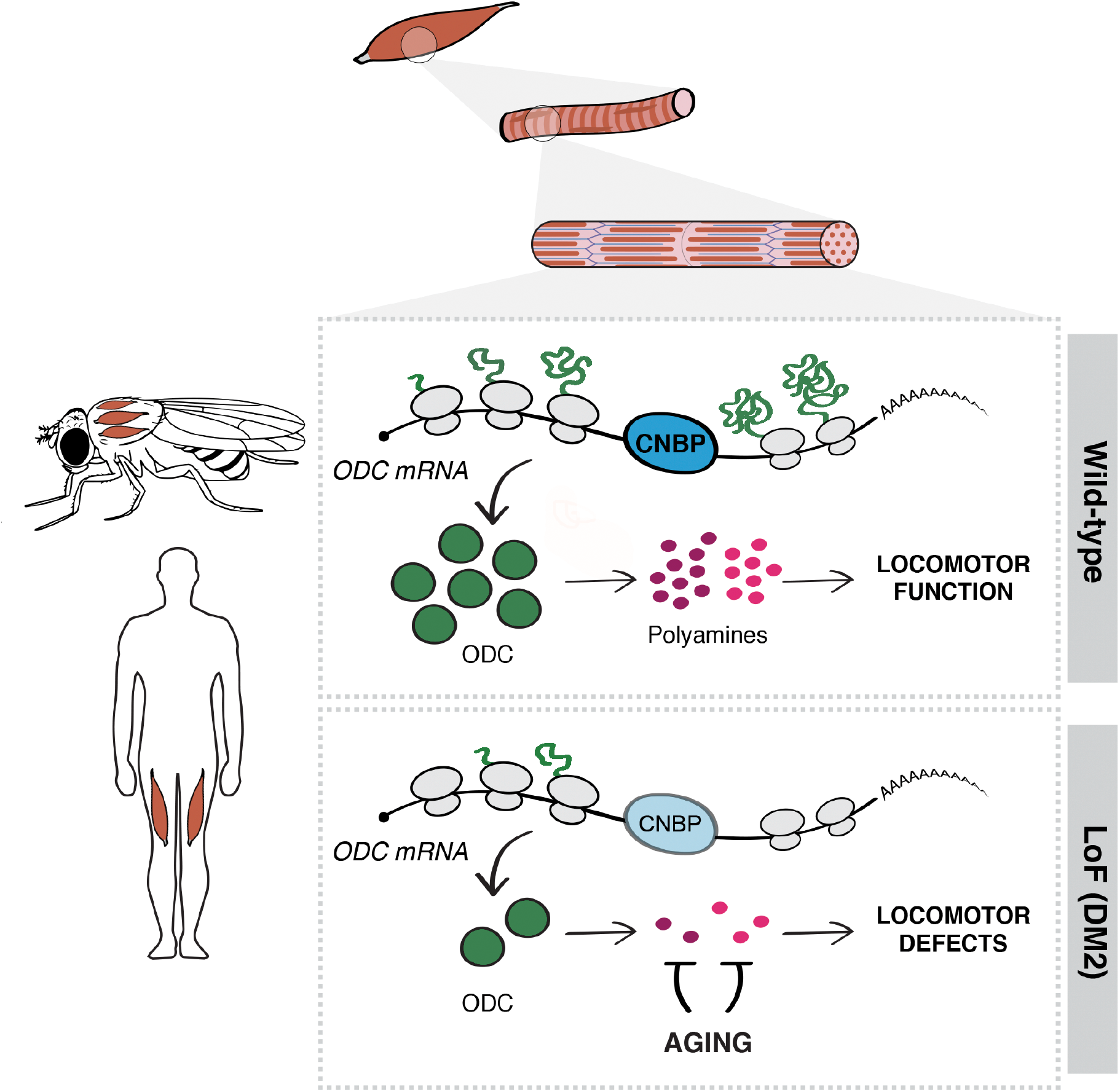

CNBP controls muscle function by regulating the polyamine metabolism

- Lack of dCNBP impairs locomotor function through ODC-polyamine downregulation
- dCNBP binds dOdc mRNA and regulates its translation
- Polyamine supplementation or dOdc1 reconstitution rescues locomotor defects
- CNBP-ODC-polyamine levels are reduced in muscle of DM2 patients

## Introduction

Myotonic dystrophy (DM) is the most common inherited muscle dystrophy in adults and comprises two genetically distinct forms: DM type 1 (DM1, Steinert’ disease, OMIM 160900), caused by an expansion of CTG repeats in the 3′ untranslated region of the DM protein kinase (*DMPK*) gene (Brook *et al*, 1992) and DM type 2 (DM2, OMIM 602668), due to the expansion of CCTG repeats in the first intron of the cellular nucleic acid-binding protein (*CNBP*) gene, also named *ZNF9* (Liquori *et al*, 2001). Both DM1 and DM2 display a multisystemic involvement of the skeletal muscle, heart, eye, brain, endocrine system and smooth muscle. The similarities in the clinical features have led to the hypothesis of a common pathogenic mechanism, represented by toxic gain of function of RNAs transcribed from alleles containing expanded CUG or CCUG repeats. These RNAs are ubiquitously transcribed, folded into hairpin structures and accumulated in nuclear foci, affecting the function of RNA-binding proteins such as the muscleblind-like proteins (MBNL1-3) and CUG-binding protein 1 (CUG-BP1) that regulates alternative splicing (Kanadia et al., 2006; Mohan et al., 2014). Recently, the involvement of additional non-mutually exclusive mechanisms, such as bi-directional antisense transcription, alteration of microRNA expression and non-ATG-mediated translation (RAN) have been demonstrated (Cho et al., 2005; Juźwik et al., 2019; Moseley et al., 2006; Nguyen et al., 2019; Perbellini et al., 2011). In particular, ectopic RAN translation has been reported in several degenerative diseases caused by microsatellite expansions such as SCA8, DM1, fragile X-associated tremor ataxia syndrome (FXTAS), C9ORF72 amyotrophic lateral sclerosis/frontotemporal dementia (ALS/FTD), Fuchs endothelial corneal dystrophy, SCA31, Huntington disease and recently in DM2 (Zu et al., 2011; Zu et al., 2013; Mori et al., 2013; Ash et al., 2013; Todd et al., 2013; Bañez-Coronel et al., 2015; Zu et al., 2017; Ishiguro et al., 2017; Soragni et al., 2018; Nguyen et al., 2019).

The presence of a repeat expansion might also lead to loss of function of the protein encoded by the affected mRNA. Haploinsufficiency of *CNBP* gene, also referred as *zinc finger protein 9* gene (*ZNF9*), resulting from the nuclear sequestration and/or altered processing of expanded pre-mRNAs, has been proposed to play an important role in the pathogenesis of DM2. In mice, heterozygous deletion of one *CNBP* allele causes a phenotype reminiscent of DM2 myopathy that gets worse with age, while homozygous deletion causes muscle atrophy and severe locomotor dysfunction already in young mice (Chen et al., 2007; Wei et al., 2018). Studies on muscle tissues or myoblasts from DM2 patients provided controversial results: some studies found normal *CNBP m*RNA and protein levels in muscle tissues (Raheem *et al*, 2010), while recent findings documented reduced levels and altered splicing of *CNBP* pre-mRNA, with corresponding decreased protein levels in muscle tissues (Huichalaf *et al*, 2009; Raheem *et al*, 2010; Salisbury *et al*, 2009; Schneider-Gold & Timchenko, 2010; Wei *et al*, 2018). The hypothesis that CNBP protein deficiency plays a key role in DM2 pathogenesis implies that perturbation of CNBP function contributes to this disease.

CNBP is a highly conserved ssDNA-binding protein of 19 kDa (Calcaterra *et al*, 2010) involved in the control of both transcription, by binding to ssDNA and unfolding G-quadruplex DNAs (G4-DNAs) in the nuclei, and translation, by binding to mRNA and unfolding G4 related structures in the cytosol (Armas *et al*, 2008; Benhalevy D, 2017; David *et al*, 2019; Huichalaf *et al*, 2009; Iadevaia *et al*, 2008).

Additionally, CNBP promotes IRES-dependent translation of the ornithine decarboxylase (ODC) mRNA working as IRES-transacting factor (ITAF; Gerbasi and Link, 2007; Sammons et al., 2011). In our previous studies, we found that CNBP promotes IRES-mediated translation of ODC and polyamine metabolism in neurons and that this mechanism is aberrantly activated in the medulloblastoma (D’Amico et al., 2015). Hence, these studies highlighted the ability of CNBP to control polyamine metabolism and illustrated the consequence of an aberrant function of this molecular regulatory mechanism in human disease.

Polyamines (putrescine, spermidine and spermine) are ubiquitous positively charged aliphatic amines that control key aspects of cell biology, such as cell growth, cell death, replication, translation, differentiation and autophagy (Casero *et al*, 2018; Coni *et al*, 2019; Wallace, 2000). Polyamine metabolism starts from the decarboxylation of ornithine into putrescine, then putrescine is converted into spermidine, which is in turn transformed into spermine (Casero *et al*, 2018; Wallace *et al*, 2003). Because of their critical role, the intracellular concentration of polyamines is tightly regulated. Conversion of ornithine into putrescine, catalyzed by ODC, an enzyme with an evolutionarily conserved function, represents the rate limiting step (Sammons *et al*, 2011; Wallace *et al*, 2003). Indeed, the intracellular levels of ODC are promptly adjusted to the cellular needs thanks to different mechanisms affecting its protein stability, transcription and translation (Pegg, 2006). Alterations of polyamine content are found in different pathophysiological conditions such as cancer, degenerative disorders and aging (Casero et al., 2018), although their specific role in muscle disorders has not been fully characterized yet.

Given the conservation of CNBP primary structure and function between *Drosophila melanogaster* and vertebrates/humans (Antonucci *et al*, 2014; D’Amico *et al*, 2015) in this work we investigate the effect of *CNBP* loss of function. We show that dCNBP depletion in muscles reduces fly viability and causes a robust locomotor defect. Furthermore, we demonstrate as dCNBP directly affects polyamine metabolism by regulating dOdc mRNA translation and, notably, that the restoration of proper polyamine content rescues muscle function.

## Results

### *CNBP* is essential for fly locomotion

To explore the role of CNBP we conducted RNAi-mediated knockdown experiments of *Drosophila dCNBP* gene. As previously shown, the expression of two copies of the RNAi construct (2XUAS*dCNBP*^*RNAi*^) under the control of ubiquitous promoters resulted in embryonic or larval lethality (Antonucci *et al*, 2014). Thus, to address the *in vivo* function of *dCNBP*, we drove the expression of 2XUAS*dCNBP*^*RNAi*^ using tissue-specific GAL4 drivers (Table1). We did not observe any effect on viability or fly locomotion activity even when *dCNBP* was efficiently silenced in neurons, motor-neurons or glia (Table1). Conversely, the expression of 2XUAS*dCNBP*^*RNAi*^ driven by the multi-tissue *69B-GAL4* or the muscle-specific *Myocyte enhancer factor 2 (Mef2)-GAL4* drivers caused early lethality at 29°C (Table1). At 25 or 18°C, lethality was reduced, allowing phenotypic analysis of adult “escapers”. Adult flies carrying 2XUAS-*dCNBP*^*RNAi*^ and either *Mef2-GAL4* or *69B-GAL4* showed strong locomotion defects (Figure 1A).

**Table 1.**
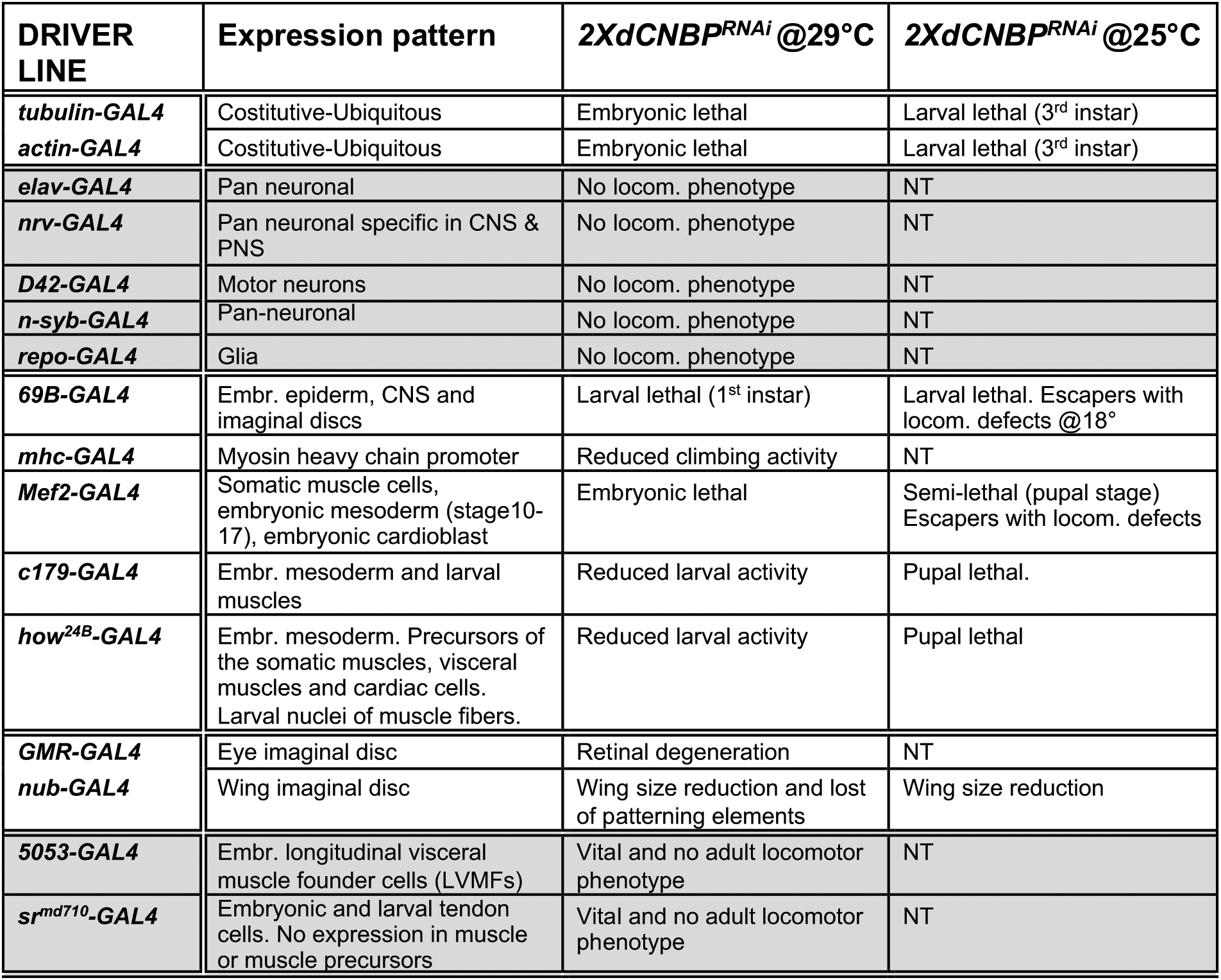
GAL4 lines expression patterns and phenotypic effects of *dCNBP* tissue-specific knockdown. In grey are represented GAL4 lines that do not cause any phenotypical effect when crossed with 2X*UASdCNBP*^*RNAi*^

**Figure 1.**
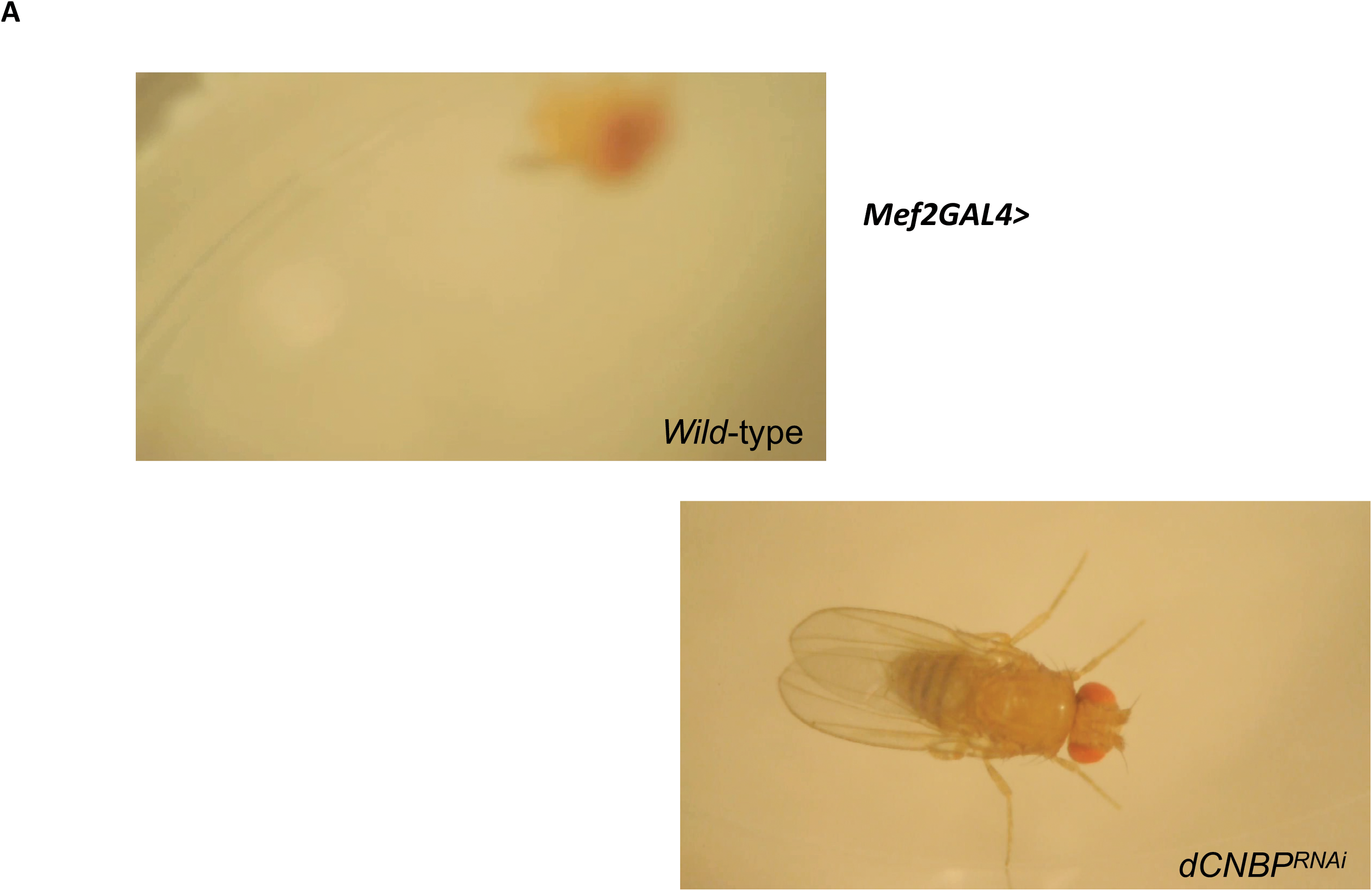
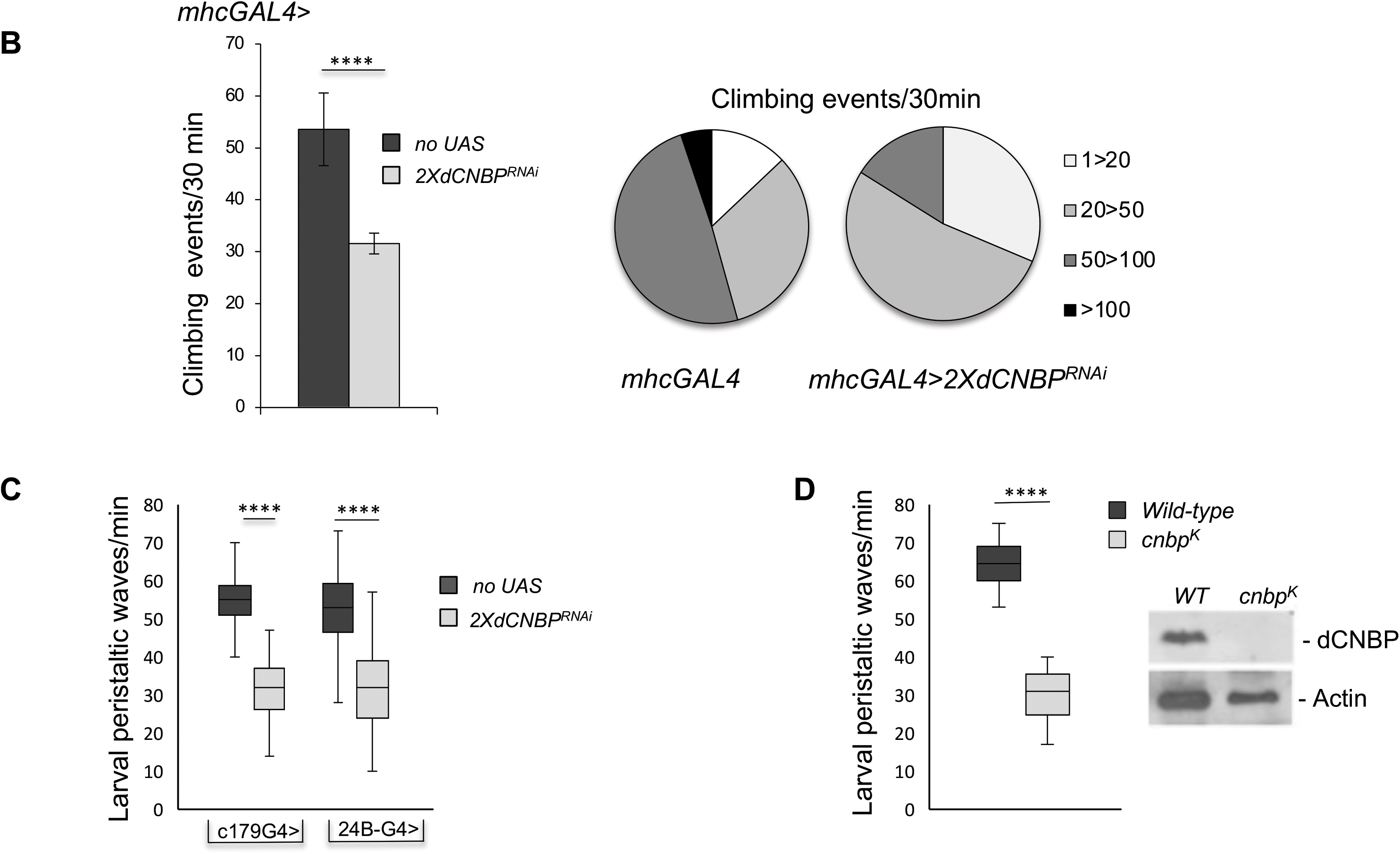
Specific *dCNBP* depletion in muscle tissues results in locomotor defects. A. Locomotion activity in escapers adult flies expressing 2X*UAS-dCNBP*^*RNAi*^ driven by the *mef2*-*GAL4* at 25°C. B. Climbing activity defects in adult flies expressing 2X*UAS-dCNBP*^*RNAi*^ driven by the *mhc*-*GAL4* at 29°C. The locomotion activity was measured by the *Drosophila* activity monitoring system (DAM), as the number of climbing events in 30 minutes. ≥ 80 males tested for each genotype. On the left, climbing performance of control flies (dark grey column) or *dCNBP* depleted flies (light grey column) 7 day after eclosion, represented as the average of climbing events (CEs) in 30 min (Error bars represent SEM; **** *P <* 0.0001, Mann-Whitney-Wilcoxon test). On the right, quantitative grouping of climbing performances in 4 different classes. Classes 1-20 (white area) and 20-50 (light grey area) CEs are highly represented in RNAi flies (*mhcGAL4>2XUASdCNBP*^*RNAi*^), while classes 50-100 CEs (dark grey area) are more frequent in control flies (*mhcGAL4*). Only control flies have the ability to perform more than 100 CEs in 30 min (black area). C. Box plot representation of the distribution of peristaltic contraction rates performed in 1min by control (*no UAS*; dark grey box) or 2X*UAS-dCNBP*^*RNAi*^ (light grey box) third instar larvae under the control of either *c179GAL4* or *24BGAL4* driver at 25°C. (**** P<0.0001, t-test). ≥ 10 larvae tested for each genotype in at least three independent experiments. D. Box plot representation of the distribution of peristaltic contraction rates performed by *wild-type* control (dark grey box) or *cnbp*^*k*^ mutant (light grey box) second instar larvae in 1 minute. ≥ 10 larvae tested for each genotype. (*P <* 0.0001, Mann-Whitney-Wilcoxon test) and levels of dCNBP protein in extract obtained from *cnbp*^*k*^ mutant second instar larvae. Actin, loading control. In (C and D) the line inside the box indicates the median for each genotype and box boundaries represent the first and third quartiles; whiskers are min and max in the 1.5 interquartile range.

https://youtu.be/caPRMRqHmw8 (movie1: *Mef2-GAL4*>2X*dCNBP*^*RNAi*^)

https://youtu.be/UAXZJz3bWwU (movie2: *Mef2-GAL4*>2X*dCNBP*^*RNAi*^)

https://youtu.be/7mP07nfMu6E (movie3: *Mef2-GAL4*>*Or-R*)

To study the effects of *CNBP* knockdown in differentiated muscle, we utilized an *Mhc*-GAL4 driver, which induces the expression of transgenes later during muscle development, compared to *Mef2-GAL4*. Although these flies were viable, they showed defective locomotion, indicating that the integrity of *dCNBP* expression is required for locomotor activity also at later stages of muscle development. Wild-type flies usually display a strong negative geotactic response: when tapped to the bottom of a vial they rapidly run to the top. As they get older or manifest locomotion dysfunction, flies no longer climb to the top of the vial, but make short abortive climbs and fall back to the bottom. The climbing activity of *mhcGAL4>*2XUAS*dCNBP*^*RNAi*^ flies was measured using the *Drosophila* activity monitoring system (DAM, TriKinetics Inc Waltham, MA USA; see Materials and Methods), which allows to quantify fly locomotion capabilities based on their negative geotactic response. As shown in Figure 1B, *mhcGAL4>*2XUAS*dCNBP*^*RNAi*^ flies exhibited a strong reduction (∼ 50%) in the number of climbing events performed in 30 minutes compared to control flies.

We also investigated the consequences of *dCNBP* silencing in the embryonic mesoderm and larval muscles. RNAi constructs expressed under the control of the c*179-* and *how*^*24B*^*-GAL4* drivers (Table1) caused late pupal lethality at both 25 and 29°C (Table1). We thus analyzed the activity of RNAi-expressing larvae by measuring the numbers of peristaltic waves performed in 1 minute. As shown in Figure 1C, both drivers caused a significant reduction of peristaltic waves compared to controls.

To further validate our findings and exclude that the phenotype could be linked to non-specific RNAi effects, we turned to an on-locus loss of function allele (dubbed *dCNBP*^*k*^). *dCNBP*^*k*^ carries a P element insertion in the *dCNBP* locus (*CG3800*) causing lethality when homozygous (larvae die at the 2^nd^ instar). *dCNBP*^*k*^ mutant larvae were examined for the expression of the dCNBP protein by immunoblotting and for their locomotor phenotype. We found that in these larvae the *dCNBP* protein product is completely absent compared to wild type (Figure 1D right). We analyzed the locomotion activity of *dCNBP*^*k*^ mutant larvae by measuring the numbers of peristaltic waves/min. As shown in Figure 1D, we found that, consistent with the RNAi data, *dCNBP* mutant larvae displayed a significant reduction of peristaltic waves compared to wild type controls.

To further confirm that locomotor defects are a specific consequence of dCNBP depletion, we generated transgenic flies bearing an RNAi-resistant cDNA (UAS*dCNBP-3HAres)* which contains appropriate synonymous substitutions in the *dCNBP* coding sequence to be resistant to RNAi-mediated degradation. Expression of this construct under the control of the *c179*-*GAL4* driver rescued larval dCNBP loss-dependent locomotor phenotype (Figure 2), confirming that this phenotype is specifically caused by *dCNBP* depletion.

**Figure 2.**
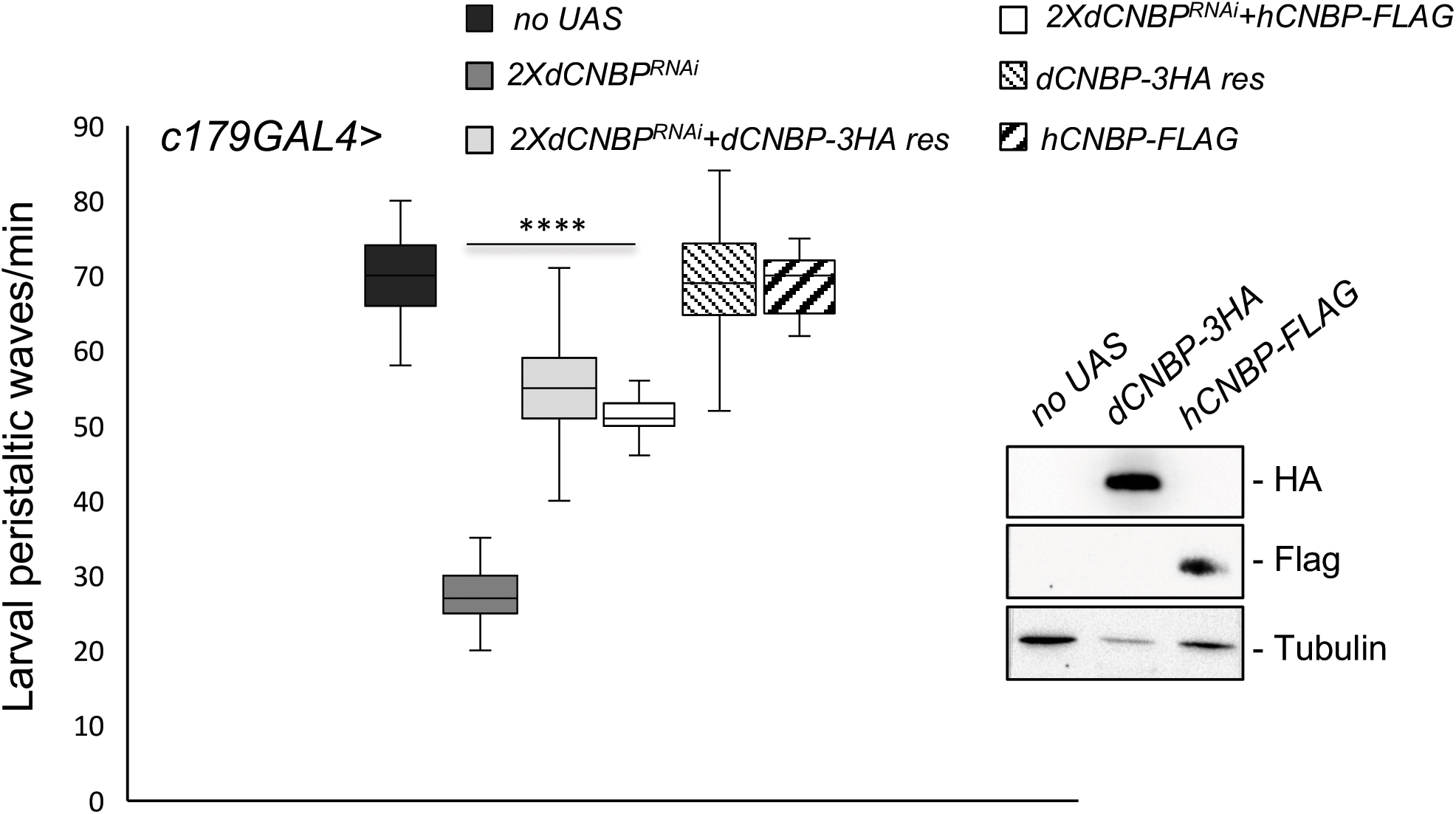
CNBP overexpression rescues the locomotion phenotype induced by muscular *dCNBP* depletion. *dCNBP* knockdown in embryonic mesoderm causes a significant reduction of larval peristaltic waves rescued by the expression of either *dCNBP* or *hCNBP* transgenes (25°C). Box plot representation of the distribution of peristaltic contraction rates performed by third instar larvae of the following genotypes: only *c179GAL4* driver (no UAS; black box), *c179GAL4*> 2X*UASdCNBP*^*RNAi*^ (white box), *c179GAL4>*2X*UASdCNBP*^*RNAi*^ + *dCNBP-HA-res* (a *dCNBP-3HA* transgene resistant to 2X*UASdCNBP*-induced RNAi, dark grey box), *c179GAL4*>2X*UASdCNBP*^*RNAi*^ + *hCNBP-FLAG* (a transgene overexpressing the human CNBP counterpart, light grey box). The line inside the box indicates the median for each genotype and box boundaries represent the first and third quartiles; whiskers are min and max in the 1.5 interquartile range (****P<0.0001, Kruskal-Wallis with post-hoc Dunn’s test). ≥ 10 larvae tested for each genotype in at least three independent experiments. Transgenes expression levels were analyzed by western blotting (right; Tub: tubulin, loading control).

We next investigated whether the human CNBP orthologue (hCNBP) could functionally rescue the prominent *dCNBP* phenotype, by expressing a *UAS hCNBP-FLAG* construct in the *Drosophila* muscle by using the *c179-GAL4* driver. As shown in Figure 2, hCNBP reversed the locomotion defects of the *dCNBP*^*RNAi*^ depleted larvae (Figure 2), indicating that hCNBP locomotor function is evolutionally conserved from fly to human.

### The ODC-polyamine pathway is involved in CNBP loss-of-function locomotor phenotype and is downregulated in human cells from DM2 patients

Having found that the absence of dCNBP in muscle tissues causes significant locomotor defects, we next sought to identify the CNBP-regulated mechanisms responsible for the observed phenotype. Since we had previously found that mammalian CNBP regulates polyamine metabolism by affecting translation of ODC in cancer cells (D’Amico et al., 2015; Sammons et al., 2011; Benhalevy et al., 2017), we asked if a similar mechanism could play a role in this context.

We performed Western blotting analysis of *dCNBP*-deficient larvae and observed that the levels of ODC are significantly reduced in larvae lacking dCNBP (in both *dCNBP* RNAi-expressing and *cnbp*^*k*^ mutant larvae, Figure 3A and Figure 3–figure supplement 1, respectively), compared to wild type controls. Accordingly, the content of putrescine, the downstream product of ODC enzymatic activity, was strongly reduced (Figure 3B and Figure 3–figure supplement 1), confirming that dCNBP regulates also ODC and polyamine levels in flies.

**Figure 3.**
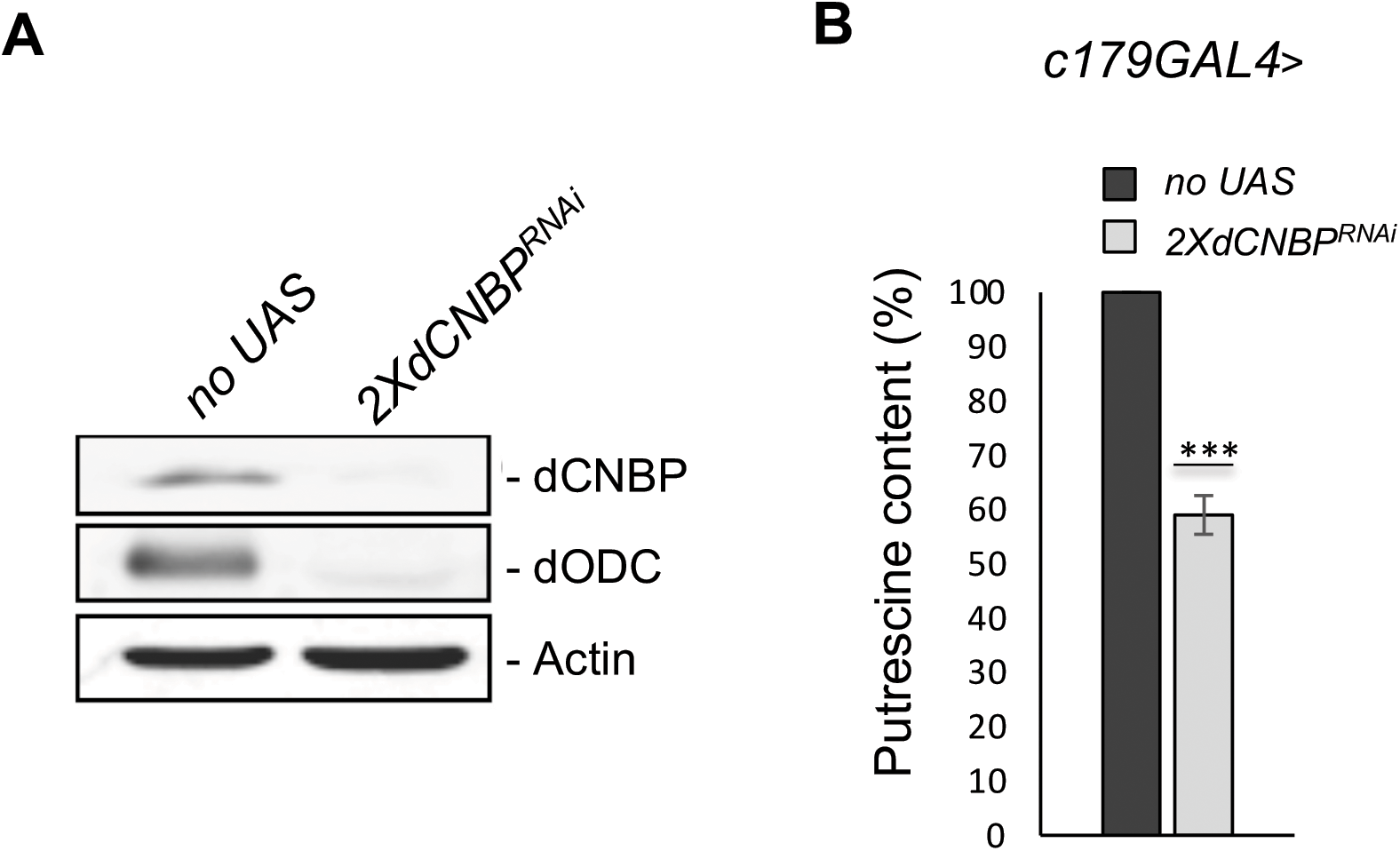
dCNBP regulates the ODC/polyamine axis. A. Levels of both ODC protein and putrescine are significantly reduced in *dCNBP*-depleted larvae compared to wild type controls. B. Immunoblot showing the levels of both dCNBP and dODC protein in extract obtained from*tubGAL4>*2X*UASdCNBP*^*RNAi*^ third instar larvae compare to control (No UAS). Actin, loading control. C. Columns represent the fold difference of putrescine content in third instar larvae bearing the *c179GAL4* driver alone (no UAS; black column) or in combination with double copy dCNBP RNAi-expressing larvae (2X*UASdCNBP*^*RNAi*^; light grey column). Error bars represent SEM; ***p>0.001, ** p>0.002, in unpaired t-test. A pool of 10 larvae has been tested for each genotype in three independent experiments.

To determine whether the locomotion defect caused by CNBP deficiency is linked to this mechanism, we analyzed the consequences of *dOdc* loss of function. We achieved a RNAi-mediated repression of both fly *Odc* genes, singularly or together (*dOdc1* and *dOdc2*; Rom and Kahana, 1993), by crossing RNAi lines (VDRC 30039 and 104597) to the *c179GAL4* driver (Figure 4A). *Odc* depletion caused adult lethality while in larvae displayed a significant impairment of peristaltic waves linked to larval locomotion activity and reduction of putrescine levels (Figure 4–figure supplement 1). Similarly, d*Odc1*^*MI10996*^ (Bloomington #56103) mutant flies displayed significant locomotor defects associated with polyamine decrease (Figure 4B). Finally, we observed that feeding of wild type flies with DFMO, an irreversible ODC inhibitor, caused a significant reduction of larval motility (Figure 4C) demonstrating that genetic or pharmacological inhibition of dOdc strongly phenocopies dCNBP downregulation effects. To determine if the ODC-polyamine axis is impaired also in the human disease, we studied muscle biopsies obtained from DM2 patients compared to those from healthy individuals. Remarkably, immunoblot analysis showed that the levels of both CNBP and ODC were reduced in DM2 patients compared to controls (Figure 5A). Consistently, we found that the content of the ODC metabolite putrescine was also significantly reduced in DM2 patients, thus indicating that polyamine synthesis might indeed be downregulated in these patients (Figure 5B).

**Figure 4.**
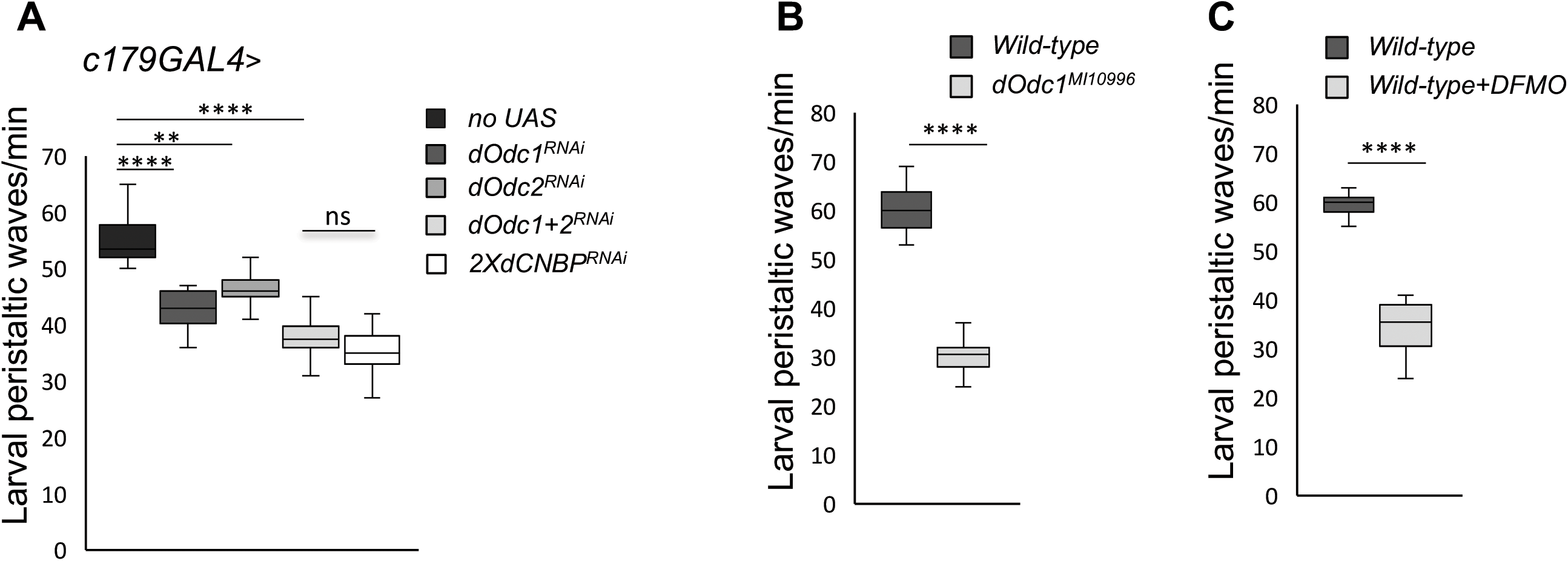
ODC depletion phenocopies the *dCNBP* locomotor defects. Box plot representation of the distribution of peristaltic contraction rates performed by the reported genotypes in one minute. A. In the legenda UAS in transgenic RNAi lines is omitted for simplicity B. W*ild type* controls (Oregon R; dark grey box) and *dOdc1*^*MI10996*^ mutant larvae (light grey box); C. W*ild type* controls fed with standard fly food (*Oregon R* dark grey box) or after DFMO treatment (5 mM/day; light grey box). D. The line inside the box indicates the median for each genotype and box boundaries represent the first and third quartiles; whiskers are min and max in the 1.5 interquartile range (**P<0.001; ****P<0.0001; ns, not significative, Kruskal-Wallis with post-hoc Dunn’s test for multiple comparison or Mann-Whitney-Wilcoxon test for). ≥ 10 larvae tested for each genotype in at least three independent experiments.

**Figure 5.**
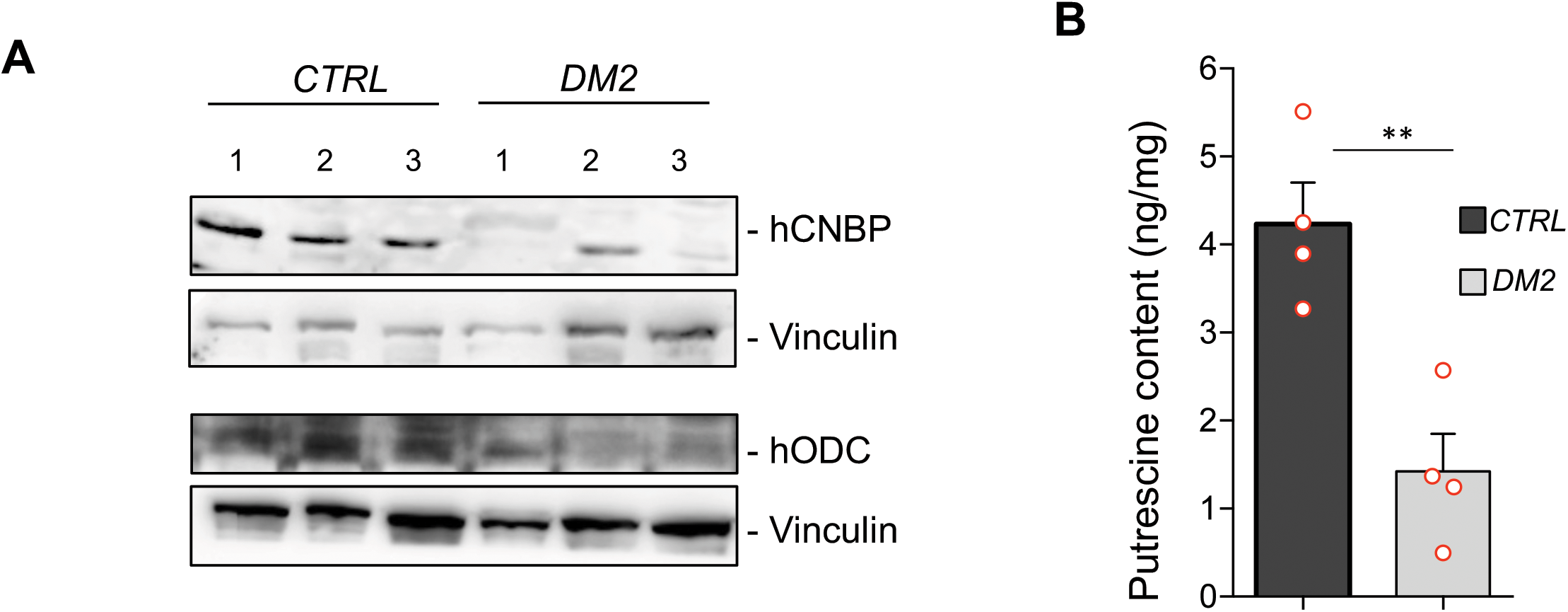
Polyamine metabolism is impaired also in DM2 muscles. CNBP and ODC content correlates with polyamine levels in DM2 muscle cells. A. Immunoblot showing the levels of both hCNBP and hODC proteins in DM2 or control muscle cells. Vinculin, loading control. B. Columns represent putrescine content in muscle cells obtained from four DM2 patients (light grey columns) or from four healthy individuals (dark grey column), expressed in ng/mg of tissue. Error bars represent SEM; ** p>0.001, in unpaired t-test.

In contrast, the levels of CNBP were not altered in a transgenic fly model of DM2 that expresses pure, uninterrupted CCUG-repeat expansions ranging from 200 to 575 repeats in length (BDSC 79583-79584-79585) and recapitulates key features of human DM2 including RNA repeat-induced toxicity, ribonuclear foci formation and changes in alternative splicing (Yu *et al*, 2015; Figure 5–figure supplement 1). These results suggest that the observed CNBP protein downregulation in DM2 patients is not a consequence of the toxic RNA accumulation, but rather a consequence of an impaired splicing process.

### dCNBP regulates dOdc translation

We next investigated the molecular mechanisms through which dCNBP controls *dOdc* expression and consequently polyamine metabolism.

We did not observe any significant reduction of *dOdc* mRNA levels in dCNBP RNAi-depleted muscles (Figure 6A), indicating that CNBP does not regulate *dOdc* mRNA synthesis or stability but rather its protein levels. In this regard, since our previous data in mammalian cells indicated that CNBP regulates IRES dependent translation of ODC (D’Amico et al., 2015), we wondered if this mechanism might also be operating in flies. To this end we cloned the 5’UTR of *dOdc1* into a bicistronic renilla-luciferase reporter vector, which allows detection of IRES activity and tested the ability of dCNBP to induce *dOdc1* IRES-mediated translation. However, ectopic expression of dCNBP, did not result in any significant change of reporter activity in mammalian cells while it significantly induced the activity of a bicistronic vector containing the 5’ UTR of human ODC, thus excluding that dCNBP could regulates *dOdc* translation through an IRES-mediated mechanism (Figure 6–figure supplement 1).

**Figure 6.**
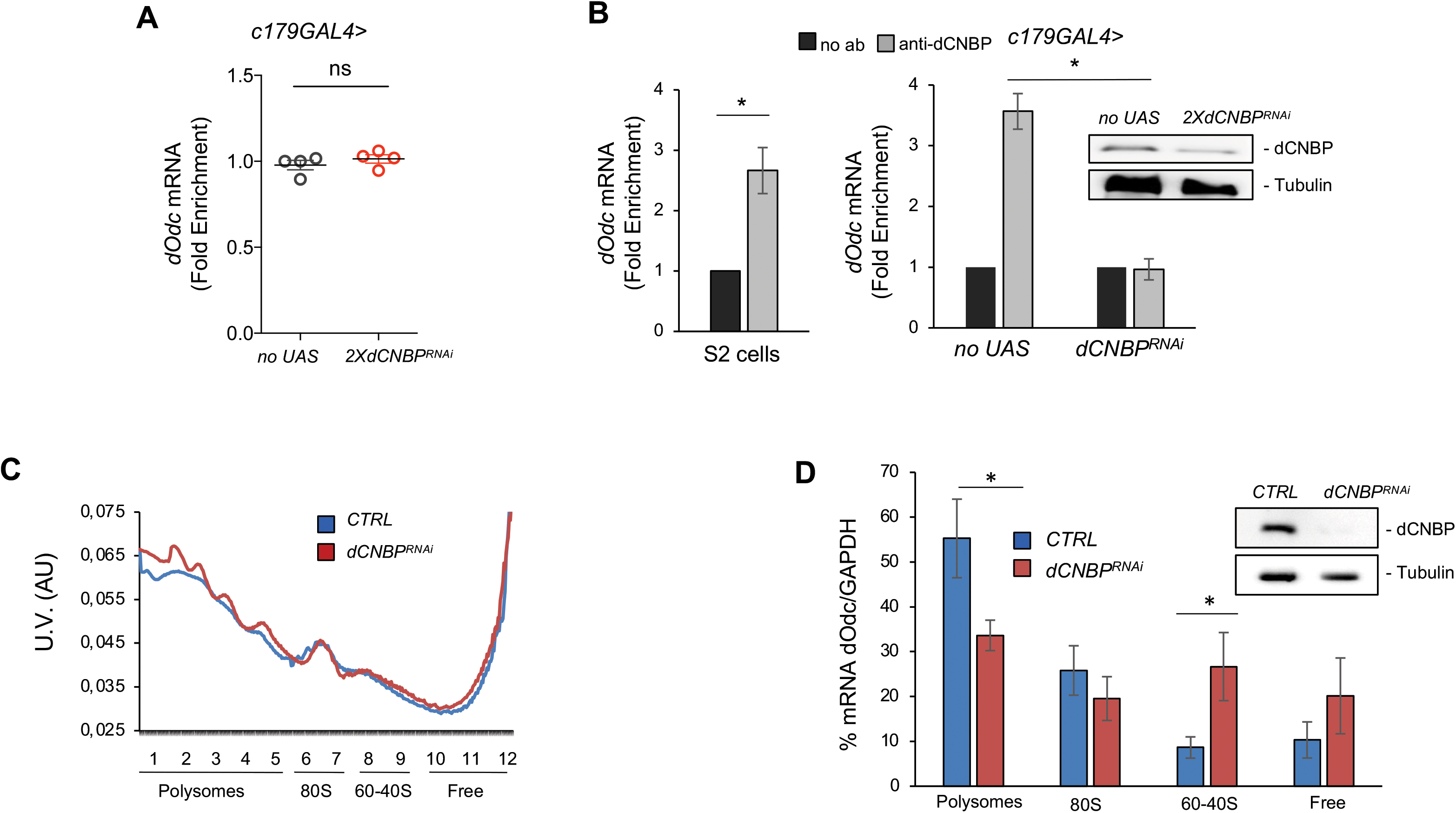
dCNBP controls polyamine metabolism through the binding and the translational control of *dOdc* mRNA. A. d*Odc1* mRNA levels (qPCR), normalized with the housekeeping *RPL11* mRNA third instar larvae bearing *C179GAL4* driver alone or in combination with UAS-*2XdCNBP*^*RNAi*^. ns: not significative in unpaired t-test. Dots correspond to four independent biological replicates; bars indicate the mean and SEM. B. CNBP binds *Odc1* mRNA. qRT-PCR analysis on mRNAs immunoprecipitated by anti-dCNBP antibody or control IgG antisera in S2 cells extracts (left), or in dCNBP-depleted (or not) larval extracts (right). The results are indicated as fold difference, relative to IgG. Error bars represent SEM of three independent biological experiments; \**P* < 0.05, in t test. The presence of dCNBP protein in C179>*2XUASdCNBP*^*RNAi*^ (or no UAS) larval carcasses was analyzed by western blotting (bottom). C. Rapresentative polysome profiles of dCNBP-deficient (in orange, *dCNBP*^*RNAi*^) or control (in blue, CTRL) S2 cells. Cytoplasmic lysates were fractionated on 15–50% sucrose gradients. D. qPCR analysis of ODC mRNA loaded in the different polysome fractions, GADPH was used to normalize the values. (\**P* < 0.05, *t*-test. Error bars represent SEM of experiments performed in quadruplicates and repeated at least three times). The presence of dCNBP protein in interferred or not S2 cells was analyzed by western blotting (right). Tubulin, loading control.

In a previous report it was shown that mammalian CNBP regulates translation of several target mRNAs via an association with G-rich recognition elements (RRE), thereby resolving their G4 stable structures and promoting translational elongation (Benhalevy et al 2017). Interestingly, one of the targets identified in that study was the mRNA of ODC and our *in silico* analysis by RBPmap (Paz *et al*, 2014; http://rbpmap.technion.ac.il) predicted the presence of several UGGAGNW motifs (the most common RRE bound by hCNBP; Figure 6–figure supplement 2) in the *Drosophila Odc1* coding sequence. Thus, we tested if dCNBP regulates translational efficiency of *dOdc1* by binding its mature mRNA. To this end, we performed RNA immunoprecipitation (RIP) assay on S2 insect cell extracts and found that CNBP efficiently binds *Odc* mRNA (Figure 6B, left). In addition, dCNBP was efficiently associated to dOdc mRNA in control (no UAS) but not in CNBP-deficient larval muscles (Figure 6B, right). The RNAi efficiency was confirmed by western blotting (Figure 6B), demonstrating the specificity of the binding. Furthermore, in a heterologous system, after transfection of a vector expressing the *dOdc* CDS, but lacking its UTRs, ODC protein synthesis was downregulated by CNBP depletion, while the mRNA levels remained unchanged (Figure 6–figure supplement 3), thus supporting the hypothesis that dCNBP regulates *dOdc* mRNA translation by acting on its coding region.

To determine whether dCNBP influences translation of *dOdc* mRNA we performed a sucrose fractionation of cytoplasmic lysates obtained from S2 cells in which *dCNBP* mRNA was knocked down or from control cells (Figure 6C and D). RNA was extracted from each fraction and analyzed by qRT-PCR (Figure 6D). The RNAi efficiency was confirmed by western blotting (Figure 6D right). In line with our assumption, in control lysates we found significant levels of *dOdc* mRNA in polysome fractions, the same where dCNBP was detected and co-purified with the ribosomal protein RpS6 (Antonucci *et al*, 2014) indicating that ODC is actively translated. In contrast, the levels of *dOdc* mRNA were strongly reduced in the polysome fraction of dCNBP-deficient S2 cells, while they were significantly increased in the non translating fractions (60-40S and free mRNA), demonstrating that dCNBP protein is required for *dOdc* mRNA loading into polysomes and therefore for its active translation (Figure 6C and D).

### Restoration of polyamine metabolism in dCNBP-deficient flies rescues locomotor phenotypes

To verify that the above-described mechanism, leading to alteration of polyamine metabolism is truly responsible of the observed locomotor phenotype, we performed rescue experiments. We first fed *dCNBP* mutant or RNAi expressing flies with putrescine dissolved into their food. As shown in Figure 7A and B, *dCNBP* mutant or RNAi-expressing larvae substantially recovered the locomotor defects after putrescine administration and, as expected, putrescine content significantly increased compared to controls reared on standard food (Figure 7–figure supplement 1A). Similar data were obtained with spermidine (Figure 7–figure supplement 1B). Then, we used the *mef2GAL4* or the *c179 GAL4* lines to drive simultaneous expression of *UAS-dOdc1* (Gupta *et al*, 2013) and 2XUAS-*dCNBP*^*RNAi*^, and found that also dOdc1 reconstitution significantly ameliorates the locomotor phenotype in *dCNBP* depleted larvae (Figure 7C). Of note, we showed that feeding with 1mM putrescine mutants for *dystrophin (Dys*^*det-1*^), a fly model of Duchenne Muscle Dystrophy (DMD), did not recover the *Dys-*dependent larval locomotor abnormalities (Figure 7D), indicating that the recovery of polyamine metabolism is specifically required for alleviating *dCNBP* loss-of-function locomotor defects.

**Figure 7.**
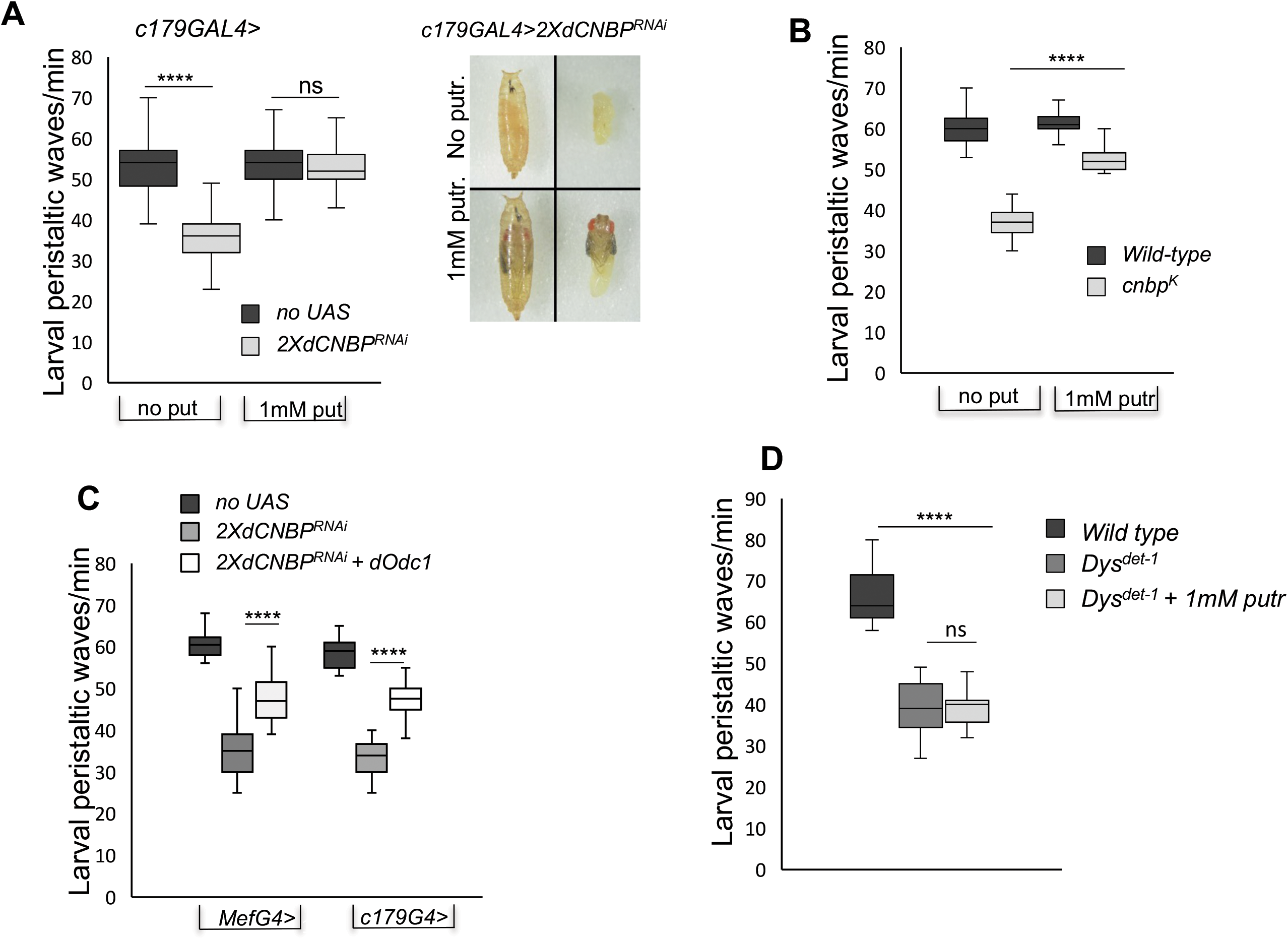
*dOdc* and polyamine are responsible of the CNBP-dependent locomotor phenotype. A-B) Rescue of locomotor phenotype in both *dCNBP* depleted larvae (A) and *dCNBP* mutant larvae (B) by 1mM putrescine feeding at 29° C. Box plot representation of the distribution of peristaltic contraction rates performed by the following genotypes: A. c*179GAL4* with or without putrescine (no UAS; dark grey boxes), *c179GAL4*>2X*UASdCNBP*^*RNAi*^ with or without 1mM putrescine (light grey boxes). Note how putrescine feeding of interfered individuals results also in a higher stage of pupal development with respect to individuals not treated (photo in A). B. *Or-R* (*wild type;* dark grey boxes) or *cnbp*^*k*^ (light grey boxes) with or without 1 mM putrescine. (**** P<0.0001; ns not significative, Kruskal-Wallis with post-hoc Dunn’s test). C. Rescue of locomotor defects in *dCNBP* depleted larvae by Odc1 overexpression under the control of either *mef* or *c179 GAL4* driver. Box plot representation of the distribution of peristaltic contraction rates performed by the following genotypes: *MefGAL4* or *c179GAL4* (no UAS; dark grey boxes), *MefGAL4* or *c179GAL4*>2X*UASdCNBP*^*RNAi*^ (light grey boxes), *MefGAL4* or *c179GAL4*>2X*UASdCNBP*^*RNAi*^ *+ UAS dOdc1* (white boxes). A-B-C) The line inside the box indicates the median for each genotype and box boundaries represent the first and third quartiles; whiskers are min and max in the 1.5 interquartile range (**** P<0.0001; ns not significative, Kruskal-Wallis with post-hoc Dunn’s test). ≥ 10 larvae tested for each genotype in at least three independent experiments. D. Mutants for *dystrophin (Dys*^*det-1*^) present larval locomotor abnormalities that cannot b*e* rescued by feding larvae with 1mM putrescine. Box plot representation of the distribution of peristaltic contraction rates performed by *Dys*^*del-1*^ mutant larvae fed with or without putrescine (dark or light grey box, respectively) with respect to wild type control (black box). The line inside the box indicates the median for each genotype and box boundaries represent the first and third quartiles; whiskers are min and max in the 1.5 interquartile range (ns, not significative, **** P<0.0001, Kruskal-Wallis with post-hoc Dunn’s test). ≥ 10 larvae tested for each genotype in at least two independent experiments.

It is known that polyamines decrease during aging in *Drosophila* (Gupta *et al*, 2013). Moreover, the decline of locomotor ability with age is common in many species of animals and muscular dystrophies gradually progress with age, along with increased muscle breakdown. Thus, we performed the DAM climbing assay with the *dCNBP* RNAi expressing adults at different time after eclosion. We found that the *dCNBP*-depleted adults showed a faster age-dependent decline of climbing ability, as 15 days aged adult flies performed a significantly lower number of climbing events/30 minutes compared to wild type control flies (Figure 7–figure supplement 2), indicating that dCNBP depletion accelerates aging-dependent locomotor decline, similar to that observed in DM patients (Mateos-Aierdi *et al*, 2015). This acceleration may well be a consequence of an aging-dependent polyamines decrease (Gupta *et al*, 2013). Together, these results, indicate, that ODC and polyamine defects are responsible of the observed CNBP loss-of-function locomotor phenotype.

## Discussion

It is widely recognized that DM1 and DM2 share many clinical features due to a common pathogenic mechanism, consisting in the toxic accumulation of RNA, resulting from the expansion of CTG triplets or CCTG quadruplets, respectively. However, the two diseases differ in some clinical manifestations, such as the preeminent involvement of proximal muscles in DM2 and of distal muscles in DM1. Therefore, it is possible that additional mechanisms contribute to the pathogenesis of the two diseases, by acting as disease modifiers. Indeed, the pathologies of repeat-expansion associated diseases are very complex, as both coding and non-coding repeat expansions may involve a combination of mechanisms, including protein loss-of-function, toxic RNA gain-of-function, and toxic protein gain-of-function. In DM2, the quadruplet expansion occurs within the first intron of the *CNBP* gene and this has given rise to the hypothesis that this genetic alteration may cause splicing defects/protein sequestration, leading to reduced CNBP protein levels. However, while some studies reported that CNBP protein levels are significantly reduced in muscle of DM2 patients, other works failed to observe such a reduction (Eisenberg *et al*, 2016, 2009; Huichalaf *et al*, 2009; Raheem *et al*, 2010; Salisbury *et al*, 2009; Schneider-Gold & Timchenko, 2010; Wei *et al*, 2018), most likely as a consequence of the limited sample sizes and the variability of the disease. Therefore, whether CNBP reduction plays a pathogenic role in DM2 is still a debated issue.

Previous studies in mice demonstrated that both heterozygous and homozygous deletion of CNBP alleles causes relevant muscle defects (Chen *et al*, 2007; Wei *et al*, 2018), suggesting a role of CNBP loss of function in the pathogenesis of the disease. In particular, while homozygous deletion of CNBP is associated to muscle atrophy and severe impairment of muscle performance at young age, the heterozygous CNBP KO mice show milder muscle dysfunctions, but develop a more pronounced locomotor phenotype at advanced age, reminiscent of DM2 disease (Wei *et al*, 2018). This latter observation is consistent with the onset of clinical manifestations in DM2 patients, which typically begins in the elderly, after the age of 60.

In the present work we have addressed for the first time the specific role of CNBP in muscle, using *Drosophila melanogaster* as a model. Using muscle-specific drivers, expressed at various stage of muscle development, we have ablated *dCNBP* gene from muscle tissues and observed severe locomotor defects that, in analogy with observations in patients and other animal models, become more pronounced with age.

CNBP deficiency is sufficient to cause this effect as evidenced by the finding that reconstitution with either dCNBP or hCNBP fully rescues the locomotor phenotype.

We have found that when *dCNBP* is knocked down at early stages of muscle development, very severe phenotypes or lethality ensue, and that knock down specifically in differentiated muscles results in robust locomotor defects. This suggests that CNBP is necessary to ensure not only proper muscle development, but also its function in the adult. Surprisingly, in contrast with studies in homozygous KO mice showing marked muscle atrophy, our morphological analysis of muscle tissues did not show significant changes upon CNBP knockdown (Figure 7–figure supplement 3). A plausible explanation for this discrepancy could be that the phenotype observed in mice is the result of the constitutive loss of CNBP in all tissues, while in our models the protein was deleted exclusively in muscle territories, likely affecting their function but not their architecture.

Mechanistically, we provide evidence that the observed phenotype is linked to the ability of CNBP to control polyamine content, by regulating ODC translation. In our previous studies, we found that in mammalian cells, CNBP binds to the 5’UTR of ODC mRNA, thereby regulating IRES-mediated translation and polyamine metabolism (D’Amico et al., 2015). Indeed, mammalian ODC mRNA has a relatively long (about 350 nts) 5’ UTR and its translation can be initiated at specific internal pyrimidine-rich sequences (Pyronnet *et al*, 2000) that were also found to bind CNBP (Gerbasi & Link, 2007). In contrast, the 5’UTR of dOdc measures only 27 nts, lacks the pyrimidine-rich sequences and does not show any IRES activity after ectopic expression of CNBP. Therefore, dCNBP does not seem to regulate translation of dOdc through an IRES-mediated mechanism.

A previous work demonstrated that CNBP facilitates translational elongation in mammalian cells, by binding G-rich motifs and resolving stable secondary structures of a number of putative transcripts, being ODC mRNA among the targets identified in that screening, although not functionally validated (Benhalevy D, 2017). These observations suggest a dual mode of CNBP regulation of ODC translation in mammals, at the level of both internal initiation and elongation across G-rich sites.

In this work we found that dCNBP binds *dOdc* mRNA and regulates its translation, likely acting at the coding region, thus supporting the conclusion that CNBP promotes *dOdc* translational elongation through the same mechanism described in mammalian cells and suggesting that the regulation of ODC by CNBP is a very important and evolutionary conserved mechanism.

Of note, in this work we have demonstrated that the locomotor defects caused by CNBP deficiency are linked to a significant decrease of polyamine content and, importantly, that the defects can be rescued by restoring dOdc expression or by polyamine supplementation. This molecular mechanism seems to be specifically linked to CNBP loss-dependent muscle dysfunction, as polyamine supplementation was unable to ameliorate the locomotor defects in a fly model of Duchenne Muscle Dystrophy. Our own data obtained from a small cohort of DM2 patients support the hypothesis that the polyamine metabolism is also altered in human DM2 muscle tissues. However, due to the heterogeneity of this disease, a study specifically addressing the representation of CNBP and polyamines, measuring their content in various muscles in a large number of patients is needed to establish more compelling evidence. Thus, whether DM2 patients may benefit from polyamine supplementation represents a crucial question opened by this work that deserves further investigation.

Interestingly, previous work demonstrated that reduced polyamine content correlates with the severity of muscle dysfunction in a mouse model of another form of human muscle dystrophy: LAMA2-Congenital Muscle Dystrophy (CMD; Kemaladewi et al., 2018). Like DM2, CMD is characterized by phenotypic variability and differentially affects specific muscle groups, possibly as a consequence of a differential expression of polyamine regulators and polyamine content.

Therefore, it is possible that CNBP also acts as a disease modifier in DM2, causing the differential distribution of polyamine content among distal and proximal muscles, which in turn sustains the clinical heterogeneity of this disease.

How polyamines affect muscle function remains to be understood. A previous study reported that supplementation of both mice and *Drosophila* diet with spermidine extends their lifespan (Eisenberg et al., 2016, 2009) and exerts protective effects on cardiac muscle of mice, by promoting cardiac autophagy, mitophagy and mitochondrial respiration (Eisenberg *et al*, 2016, 2009). Thus, it is possible that an impairment of these mechanisms in muscle may be involved in the observed phenotype of CNBP-deficient animals and possibly in DM2 patients. Moreover, polyamines are among the substances that have been reported to decline with age (Gupta *et al*, 2013; Liu *et al*, 2008) and the phenotype of CNBP-deficient animals or the clinical manifestation of DM2 patients are also correlated with the advanced age. Therefore, it is possible that polyamine may be involved, at least in part, in the age-dependent manifestations of the disease. Further studies on the role and mechanism of action of polyamines in muscle function are thus required to elucidate this critical issue.

In conclusion, we have identified an unprecedented mechanism whereby dCNBP controls muscle function by regulating the ODC/polyamine axis. This function of dCNBP we have described in *Drosophila* seems to be evolutionarily conserved in vertebrates, with relevant implications in DM2 disease.

## Author Contributions

L.C. and G.C. conceived and coordinated the project, designed experiments, analyzed the data, and wrote the paper; S.C., F.A.F. and M.Mr. designed and performed experiments and analyzed the data in both *Drosophila* and cell lines; A.M. and E.A. performed measurements and analysis of the polyamine data; B.Z., F.M. and M.Mn. performed *Drosophila* genetics and locomotor analysis. Z.I. and A.F. performed polysomal fractionation; R.B. performed cloning and part of mammalian experiments; G.S. and A.P. provided human control and DM2 patients’ muscle biopsies for IB studies; S.S. and T.M. performed fly muscles phenotypic analysis; L.C., G.C., S.C., M.Mr. and M.Mn. conceived and illustrated the graphical abstract; L.C., G.C., S.C., M.Mr., S.S. and G.S. critically revised and edited the manuscript.

## Acknowledgments

This work was supported by AFM Telethon (French Muscular Dystrophy Association, project 21025). AIRC (Associazione Italiana Ricerca Cancro IG 17575), Sapienza University grant RM1181642798C54A and Italian Ministry of Education, Universities and Research, Dipartimenti di Eccellenza-L. (232/2016). We thank Vittorio Padovano for his contribution in generating the mouse anti dCNBP antibody. We thank Gianluca Cestra for critical reading of the manuscript.

## Material and Methods

### *Drosophila* strains and rearing conditions

The *2XUAS-dCNBP*^*RNAi*^ strain used for *dCNBP* downregulation was already described in Antonucci *et al*., 2014. Essentially are transgenic flies carrying two different *UAS-dCNBP*^RNAi^ constructs (VDRC, ID 16283 and 16284) one on the X and one on the second chromosome, respectively.

The *UAS-Odc1*^*RNAi*^ and the *UAS-Odc2*^*RNAi*^ strains were also obtained from VDRC (ID 30039 and 104597) and similarly were combined to generate the strain *UAS-dOdc1-2*^*RNAi*^ bearing both constructs to downregulate both isoform at the same time.

*dCNBP*^*k*^ is one of the P element insertions in the *CG3800* locus obtained from the Kyoto DGRC (#203535). The RNAi resistant *dCNBP* gene carries synonymous substitutions in each residue of the region recognized by *UAS-dCNBP*^*RNAi*^ and was synthesized by Genewiz (SIGMA-ALDRICH). The plasmids for inducible expression of RNAi resistant *dCNBP-3HA* (abbreviated with *dCNBP-3HA-res*) was generated by cloning the 3HA epitope CDS fused in-frame with the 3′ end of the *RNAi* resistant *dCNBP* CDS into the UAS-attB vector (Genewiz, SIGMA-ALDRICH). The plasmids for inducible expression of the *human CNBP* counterpart was generated by cloning the FLAG epitope CDS fused in-frame with the 3′ end of the *hCNBP* CDS (CNBP-201 splice variant, CCDS 3056.1) into the UAS-attB vector (Genewiz, SIGMA-ALDRICH). The *dCNBP-3HA-res* or *UAS-hCNBP-FLAG* were injected in y^1^ w^67c23^; P{CaryP}attP2 embryos (BDSC Stock#8622); germline transformation was performed by Bestgene Inc. (Chino Hills, California) using standard procedures. *UAS*–*Odc-1* (Gupta *et al*., 2013). All the driver lines used have been previously described and available from the Bloomington stock center.

Spermidine or putrescine was added to normal food to a final concentration of 1 mM. For experiments, parental flies mated on either normal or Spd+ or Put+ food, and their progeny was allowed to develop on the respective food. DFMO was added to normal food to a final concentration of 5 mM/day.

### Climbing assays

The locomotion activity was measured by the *Drosophila* activity monitoring system (DAM, TriKinetics Inc Waltham, MA USA), which allows a measure of fly locomotion capabilities based on their negative geotactic response, as the number of climbing performances in 30 minutes. 10–15 age-synchronized male flies (2-3 days, 7 day or 15 days old) were gathered and placed in each monitor for each genotype for each experiment. Briefly, the *Drosophila* Activity Monitor System (Trikinetics) records activity from individual flies maintained in sealed tubes placed in activity monitors. An infrared beam directed through the midpoint of each tube measures an “activity event” each time a fly crosses the beam. The number of climbing events was scored for 30 minutes, tapping flies to the bottom every 40 seconds. Events detected over the course of each consecutive sampling interval are summed and recorded over the course of 30 minutes for each fly.

### *Drosophila* larval locomotion analyses

Larval locomotor activity was measured by counting the number of peristaltic contractions of third instar larvae performed within 1 min on the surface of a 1% agarose gel in a Petri dish; measurements were repeated five times for each larva, at least 10 larvae per genotype in each experiment.

### Immunoblot and antibodies

Protein extracts were derived from 5 third instar larvae, or cultured *Drosophila* S2 or human 293 cells, lysed in sample buffer, fractionated by SDS-PAGE and transferred to nitrocellulose membrane. Primary antibodies were: anti-CNBP goat (1:500; Abcam, Ab 48027); anti-Actin goat (1:1000; Santa Cruz, sc-1616); anti-ODC rabbit (1:500; Enzo Life Science, BML-PW8880-0100); anti-CNBP mouse (1:1000; generated by Agrobio for this work), anti-HA HRP (1:500; Santa Cruz, sc-7392), anti-GFP mouse (1:500; Santa Cruz, sc-9996); anti-FLAG HRP (1:1000; Sigma, A8592); anti-Vinculin mouse (1:1000; Santa Cruz, sc-73614); anti-Tubulin mouse (1:7000; Sigma, T-5168). As secondary antibody we used the appropriate HRP-conjugated antibody (GE Health Care) diluted 1:5000 in 5% milk/ PBS-Tween 0,1% (GE Health Care). Detection was performed by using WesternBright ECL (K-12045-D50, Advansta). Densitometric analysis was performed using ImageJ software (version 1.50i). For DM2 patient biopsies, samples were lysed in SDS urea (50mM Tris HCl ph 7.8; 2% SDS, 10% Glycerol, 1mM EDTA, 6M UREA, 50mM NaF, 5mM Na2P2O7) sonicated for 10 seconds, quantified by using a nanodrop and loaded on polyacrylamide gel. Muscle biopsies used for this study were performed primarily for diagnostic purposes, after receiving an informed consent from all patients also regarding their future use for research purposes and in agreement with the guidelines of the Ethical Committee of Our Institution.

### RNA interference in S2 cell lines

S2 cells (DGRC, RRID: CVCL_Z232; tested negative for mycoplasma) were cultured at 25°C in Schneider’s insect medium (Sigma) supplemented with 10% heat-inactivated fetal bovine serum (FBS, Gibco). RNAi treatments were carried out according to (Somma *et al*, 2008). dsRNA-treated cells were grown for 4–5 days at 25°C, and then processed for biochemical analyses. PCR products and dsRNAs were synthesized as described in (Somma *et al*, 2008). The primers used in the PCR reactions were 35 nt long and all contained a 5’ T7 RNA polymerase binding site (5’-TAATACGACTCACTATAGGGAGG-3’) joined to a gene-specific sequence.

### RNA immunoprecipitation

S2 cells were plated in 75cm^2^ flask culture dishes and 72 hours later cells were cross-linked with 1% formaldehyde solution. Pellets were lysed with FA Buffer (50 mM HEPES pH 7.5, 140 mM NaCl, 1 mM EDTA, 1% Triton X-100, 0.1% sodium deoxycholate, protease inhibitors and 50 U/ml RNase inhibitor SupeRNase, #AM2694 Thermo Fisher Scientific) and sonicated.

For *in vivo* analysis approximately 50 larval carcasses were UV crosslinked (3 × 2000 µJ/cm^2^), homogenized on ice in 1 mL RCB buffer (50 mM HEPES pH 7.4, 200 mM NaCl, 2.5 mM MgCl2, 0.1% Triton X-100, 250 mM sucrose, 1 mM DTT, 1× EDTA-free Complete Protease Inhibitors, 1 mM PMSF) supplemented with 300 U RNAseOUT (Invitrogen) and placed on ice for 30 min. The homogenate was sonicated on ice, at 80% power, five times in 20 s bursts with a 60 s rest in between using the Hielscher Ultrasonic Processor UP100H (100 W, 30 kHz) and centrifuged (16000 × g for 5 min at 4 °C). Immunoprecipitation was performed incubating the samples with anti CNBP antibody or IgG overnight. Then the samples were washed with RCB buffer four times, or with three different solutions for S2 extracts: Low salt solution: 0.1% SDS, 1% Triton X-100 2mM, EDTA 20 mM Tris-HCl pH 8, 150 mM NaCl and 0,005 U/ml SuperRNAse (Thermofisher Scientific); High salt solution: 0.1% SDS, 1% Triton X-100, 2 mM EDTA, 20 mM Tris HCl pH 8, 500 mM NaCl and 0.005 U/ml SuperRNAse; LiCl buffer solution: 0.25 M LiCl, 1% NP40, 1% sodium deoxycholate, 1 mM EDTA, 10 mM Tris-HCl pH 8 and 0.005 U/ml SuperRNAse; TE wash buffer solution: 10 mM Tris-HCl pH 8, 1 mM EDTA and 0.005 U/ml SuperRNAse, and then eluted with H2O or Elution buffer solution for S2 extracts: 1% SDS, 0.1 M NaHCO3, SuperRNase 50 U/ml. RNA was purified using Trizol reagent (15596026, Thermofisher), it was reverse transcribed and dOdc was amplified by qPCR. Results were normalized on RPL11.

### RNA extraction and quantitative PCR

Total mRNA was isolated from S2 cells or *Drosophila* larvae by using Trizol reagent (15596026, Thermofisher) according to the manufacturer’s instructions. RNA was reverse-transcribed (1 µg each experimental point) by using SensiFAST™ cDNA Synthesis Kit (BIO-65053, Bioline) and qPCR was performed using SensiFast Sybr Lo-Rox Mix (BIO-94020, Bioline). The run was performed by using the Applied Biosystems (Waltham, Massachusetts, USA) ViiA 7 Real-Time PCR System 36 instrument. The following primers were used:

dOdc Fw: TGGCAGCGATGACGTAAAGTT;

dOdc Rv: TGGTTCGGCGATTATGTGAA;

dRPL11 Fw; CCATCGGTATCTATGGTCTGGA;

dRPL 11 Rv; CATCGTATTTCTGCTGGAACCA; GFP

Fw: GCAAAGACCCCAACGAGAAG;

GFP Rv: TTCTGATAGGCAGCCTGCAC;

dGADPH Fw: CCTGGCCAAGGTCATCAATG;

dGADPH Rv: ATGACCTTGCCCACAGCCTT;

### Polysome analysis

Polysomal fractionation from S2 cells was performed as described previously (Coni *et al*, 2020), S2 cell (interfered or not) were incubated 5 minutes with100 μg/ml CHX, then washed with PBS and lysed with TNM buffer (10mM Tris-HCl pH 7.4 or 7.5, 10mM NaCl, 10mM MgCl2, 1%Triton X-100) supplemented with 10mM dithiothreitol, 100 μg/ml CHX, 1x PIC (1187358001 complete, EDTA free, Roche,), and RiboLock RNase inhibitor (EO0382, Thermo Fisher Scientific). Lysates were incubated on ice for 10 minutes and then centrifuged at 2000 rpm for 5 min. Supernatants were loaded onto 15–50% sucrose gradients and centrifuged for 120 min in a Beckman SW41 rotor at 37000 rpm at 4 °C. Fractions were automatically collected, using Biorad-BioLogic LP/2110 (Hercules) monitoring the optical density at 260 nm. RNA was extracted from each fraction by using Trizol Reagent and dOdc mRNA was amplified by RT-qPCR. GAPDH mRNA was used for normalization.

### 293T lentiviral transduction and transfection

Lentivirus production was performed as described in (D’Amico *et al*., 2015). Then human 293T cells were transduced with lentiviral particles of plkoSCR (Mission plko.1 puro; SHC002) or shCNBP human (Mission plko.1 puro TRCN0000311158, SIGMA-ALDRICH) at a MOI=5 for 72h. Then 293T cells SCR and shCNBP were transfected with plasmids encoding for dOdc and GFP for extra 24 hours, by using Dreamfect reagent according to manufacturer (DF41000 OZ, Biosciences). Cell extracts were analyzed through Western Blot and qPCR as indicated.

### Polyamine Analysis

Polyamine content was determined by gas chromatography-mass spectrometry (GC-MS) and the values were normalized by the protein concentration. A pool of 10 third instar larvae for each genotype were resuspended in 0.2 M HClO and homogenized in an ice-bath using an ultra-turrax T8 blender. The homogenized tissue was centrifuged at 13000 g for 15 min at 4°C. 0.5 ml of supernatant was spiked with internal standard 1,6-diaminohexane and adjusted to pH ≥12 with 0.5 ml of 5 M NaOH. The samples were then subjected to sequential *N*-ethoxycarbonylation and *N*-pentafluoropropionylation. For DM2 samples biopsies were also resuspended in 0.2 M HClO4 and processed as described above. GC-MS analyses were performed with an Agilent 6850A gas chromatograph coupled to a 5973N quadrupole mass selective detector (Agilent Technologies, Palo Alto, CA, USA). Chromatographic separations were carried out with an Agilent HP-5ms fused-silica capillary column. Mass spectrometric analysis was performed simultaneously in TIC (mass range scan from *m/z* 50 to 800 at a rate of 0.42 scans s–1) and SIM mode (Put, *m/z* 405; Spd, *m/z* 580, *N*1-acetyl-Spm, *m/z* 637; Spm, *m/z* 709).

### Immunostaining and confocal imaging

Larvae were dissected in ice-cold Ca^2+^-free HL3 saline and fixed in 4% formaldehyde for 10 min and washed in PBS containing 0.05% Triton X-100 (PBST) for 30 min. After washing, larval fillets were stained with phalloidin–TRITC (1:300 diluted in PBST, Sigma) for 40 min at room temperature and subsequently washed for 3x 20 min with 0,05% PBST. Larvae were mounted in Vectashield containing DAPI (Vector Laboratories).

Confocal microscopy was performed with a Leica SP8 confocal microscope (Leica Microsystems, Germany). Confocal imaging of larval fillets was done using a z step of 0.5 μm. The following objective was used: 63× 1.4 NA oil immersion for confocal imaging. All confocal images were acquired using the LCS AF software (Leica, Germany). Images from fixed samples were taken from third instar larval fillets (segment A2, muscle 6/7).

### Statistical analyses

Statistical analysis was performed using Prism 6 software (MacKiev). The Shapiro-Wilk Test was used to assess the Normal distribution of every group of different genotypes. Statistical differences for multiple comparisons were analyzed with the Kruskal-Wallis for non-parametric values or with one-way ANOVA for parametric value. The Dunn’s or the Tukey’s test was performed, respectively, as Post-Hoc Test to determine the significance between every single group. The Mann–Whitney *U* test or the t-test were used for two groups comparison of non-parametric or parametric value, respectively. A *P*< 0.01 was considered significant.

## FIGURE LEGENDS

**Figure 3–figure supplement 1.**
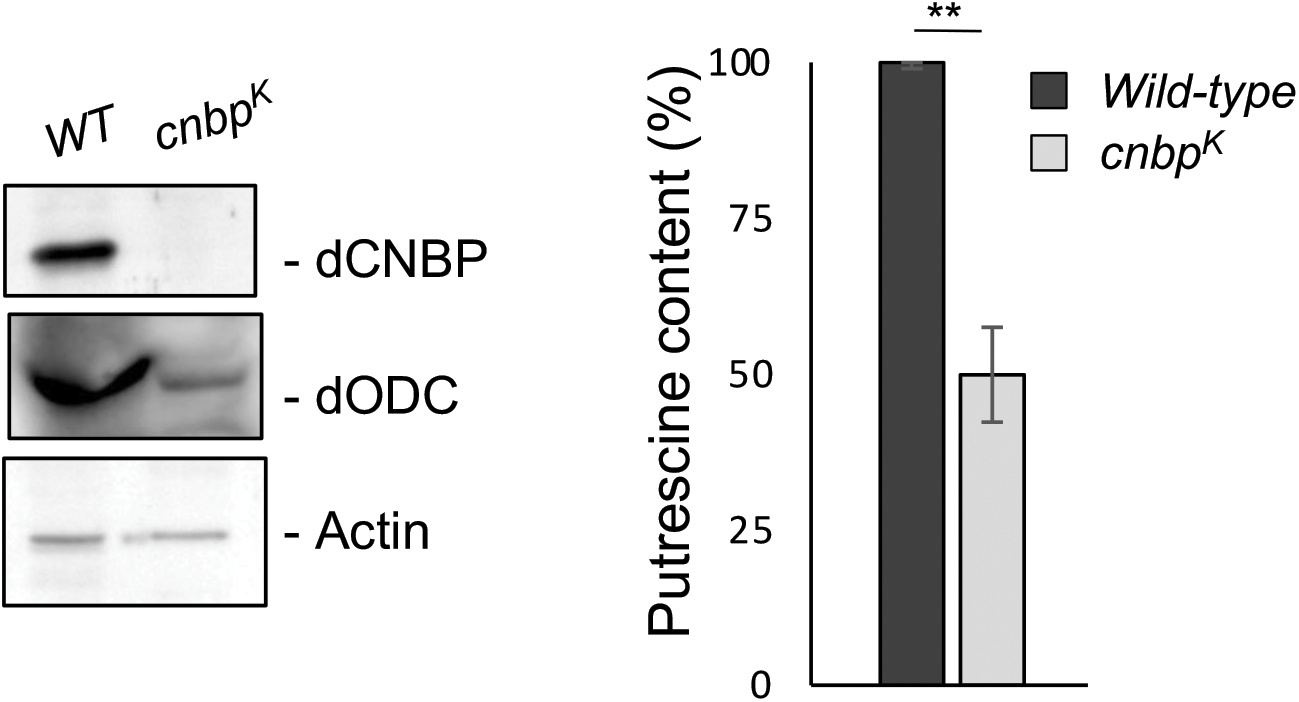
Larval locomotor defect observed in *dCNBP*^*k*^ mutants correlates with the reduction of Odc protein and polyamine levels. Levels of ODC protein and polyamine are significantly reduced in *dCNBP* mutant larvae compared to wild type controls. Left: immunoblot showing the levels of both dCNBP and dOdc protein in extract obtained from wild type (WT) and *cnbp*^*k*^ mutant second instar larvae. Actin, loading control. Right: columns represent the fold difference of putrescine content in second instar larvae of *cnbp*^*k*^ mutant (light grey column) compared to controls (wild type, dark grey column) Error bars represent SEM; ** p>0.002, t test. A pool of 10 larvae has been tested for each genotype in three independent experiments.

**Figure 4–figure supplement 1.**
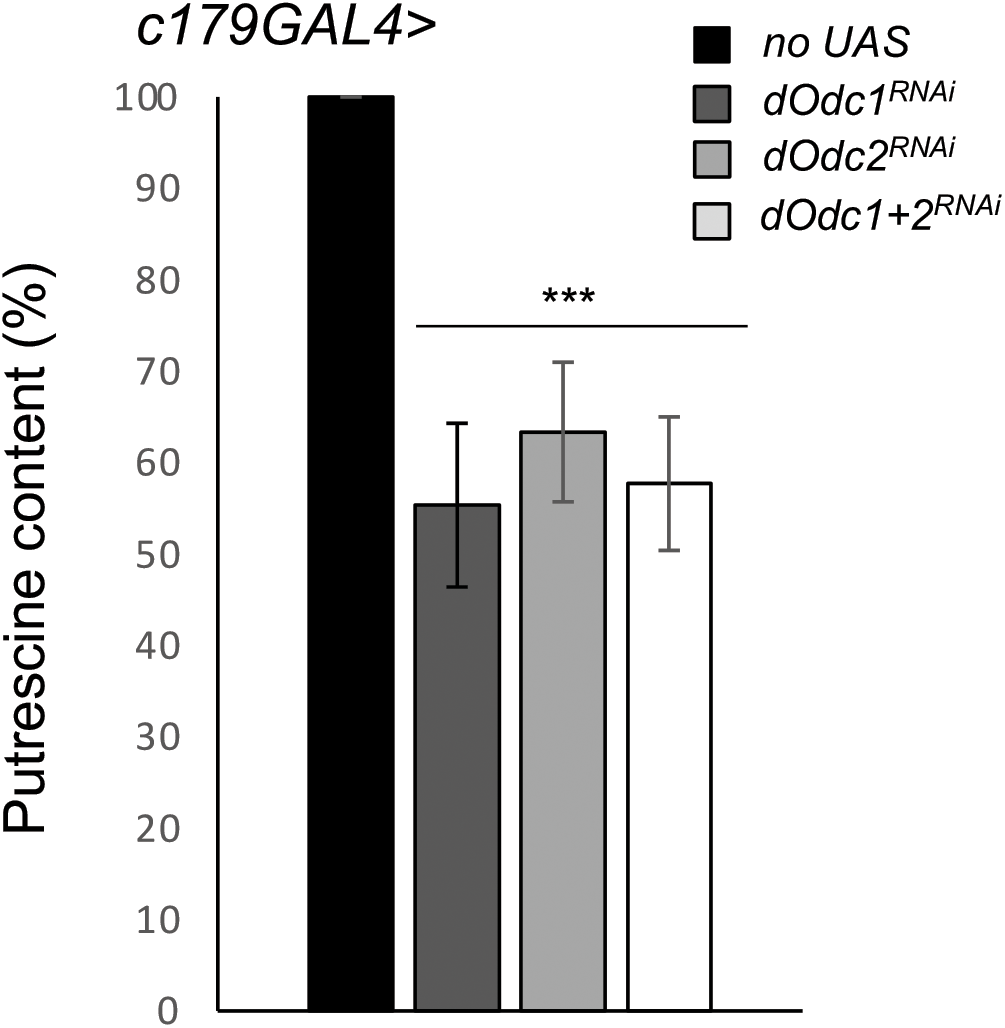
Larval locomotor defect observed as consequence of Odc depletion correlates with the reduction of polyamine levels. Columns represent the fold difference of putrescine content in third instar larvae of the following genotypes: only *c179GAL4* driver (no UAS; black column), *c179GAL4*> *UASdOdc1*^*RNAi*^ (dark grey column), *c179GAL4> UASdOdc2*^*RNAi*^ (light grey column), *c179GAL4*> *UASdOdc1*^*RNAi*^ + *UASdOdc2*^*RNAi*^ (white column). (Error bars represent SEM; ***p>0.001 respect to the no UAS control, t test). A pool of 10 larvae has been tested for each genotype in three independent experiments.

**Figure 5–figure supplement 1.**
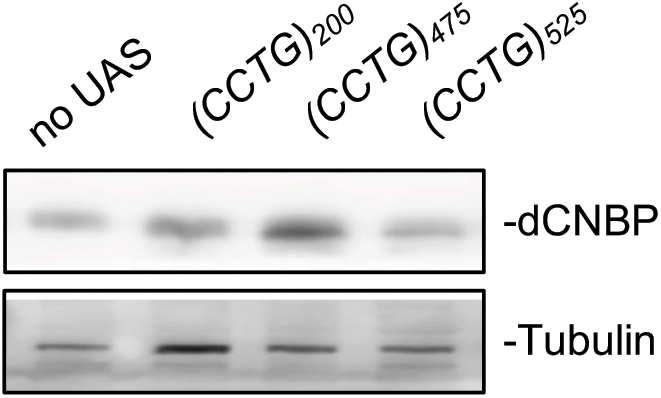
Expression levels of dCNBP are not affected by the expression of CCUG-expanded repeat RNA. Immunoblot showing the levels of dCNBP protein in larval extract obtained from controls (no UAS) or *UAS-(CCTG)200, UAS-(CCTG)475, UAS-(CCTG)525* driven by the *c179GAL4* driver. Tubulin, loading control. A pool of 10 larvae has been tested for each genotype in two independent experiments.

**Figure 6–figure supplement 1.**
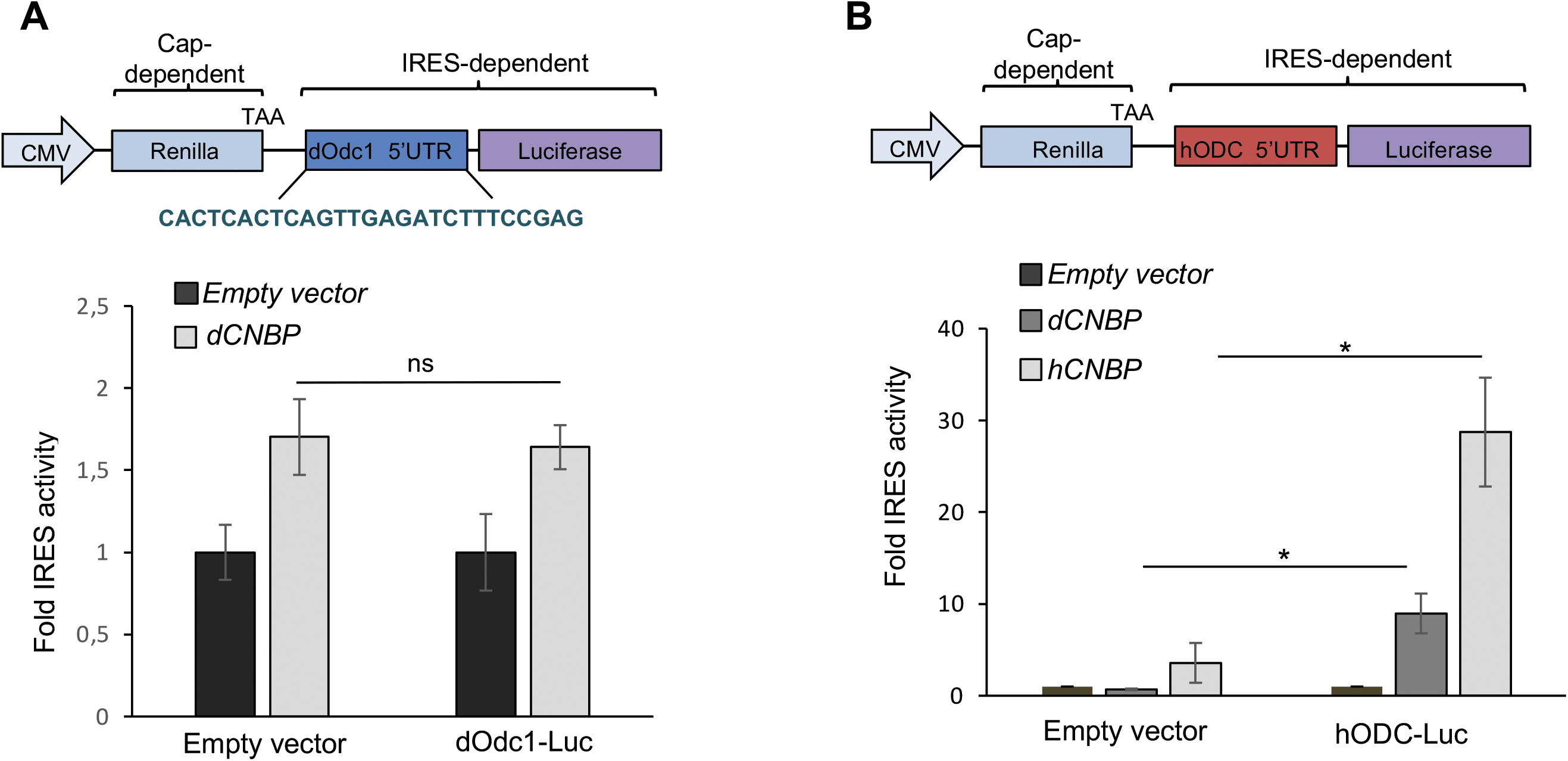
dCNBP does not control polyamine metabolism through *dOdc1* IRES-dependent translation. A. Schematic representation of the *bicistronic CMV-renilla-TAA/dOdc1-IRES-firefly luciferase* (d*Odc1-luc*) *vector* (top). The renilla ORF is translated via conventional cap-dependent mechanism, whereas translation of the luciferase ORF is controlled by the *dOdc1* 5’ UTR sequence. IRES activity of d*Odc1-luc* which is not significantly modulated by dCNBP overexpression compared to the empty vector (bottom). Columns represent the fold changes of luciferase activity, normalized to the renilla expression. Error bars represent SEM; ns: no significant with Student’s test, in three independent biological experiments B. Schematic representation of a *bicistronic CMV-renilla-TAA/hODC1-IRES-firefly luciferase* (h*ODC1-luc*) *vector* (top). IRES activity of h*ODC1-luc* which is significantly modulated by both dCNBP or hCNBP overexpression, compared to the empty vector (bottom). Columns represent the fold changes of luciferase activity, normalized to the renilla expression. Error bars represent SEM; p<0,05 with Student’s test, of three independent biological experiments.

**Figure 6–figure supplement 2.**
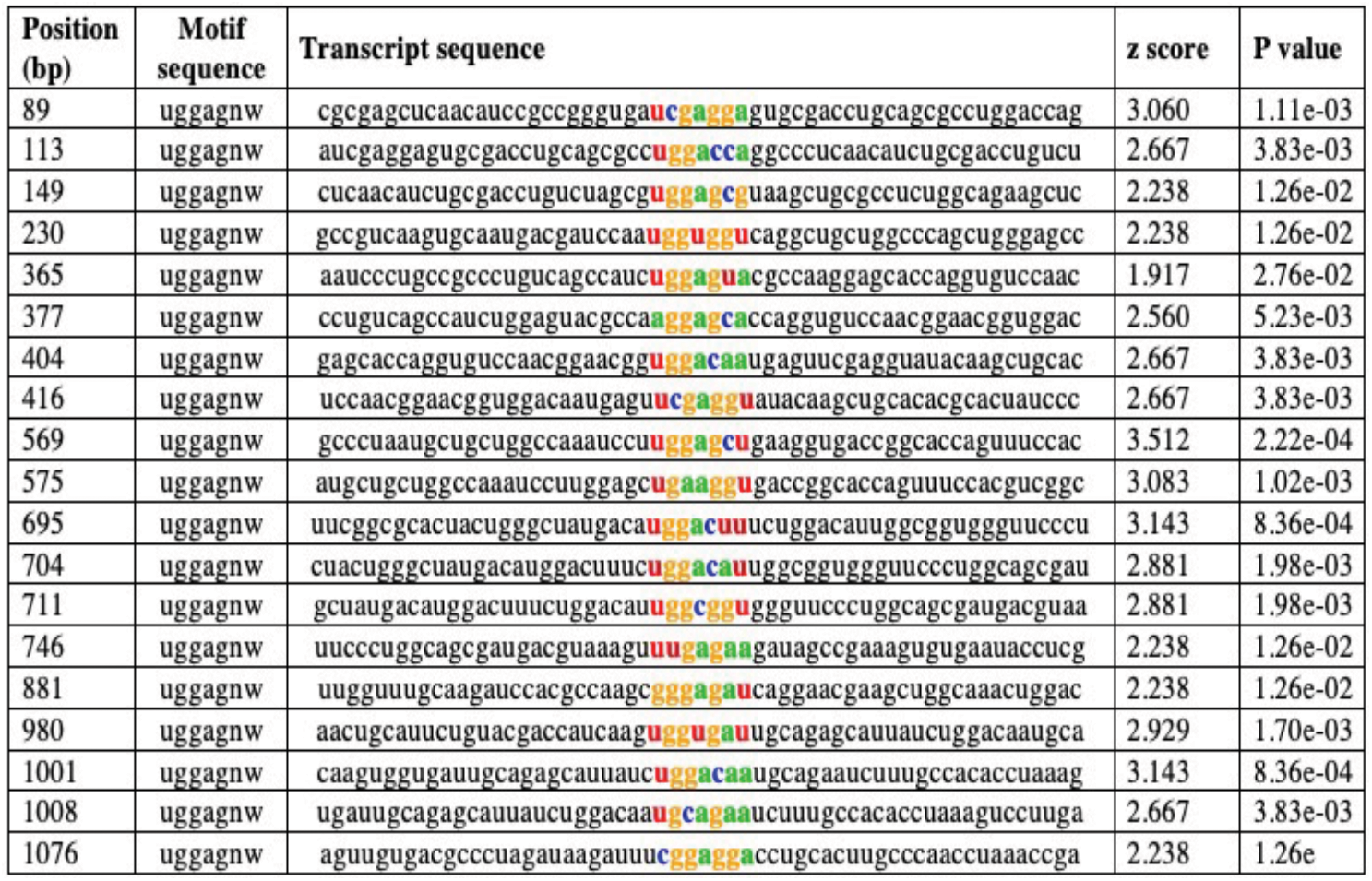
In silico prediction of putative CNBP binding sites on the *dOdc1* mRNA by RBPmap. The dOdc1 transcript (FBtr0088863) was uploaded to the RBPmap web server for mapping binding sites. As criteria *Drosophila* genome, UGGAGNW consensus motif and high stringency level were used (Paz *et al*., 2014; http://rbpmap.technion.ac.il).

**Figure 6–figure supplement 3.**
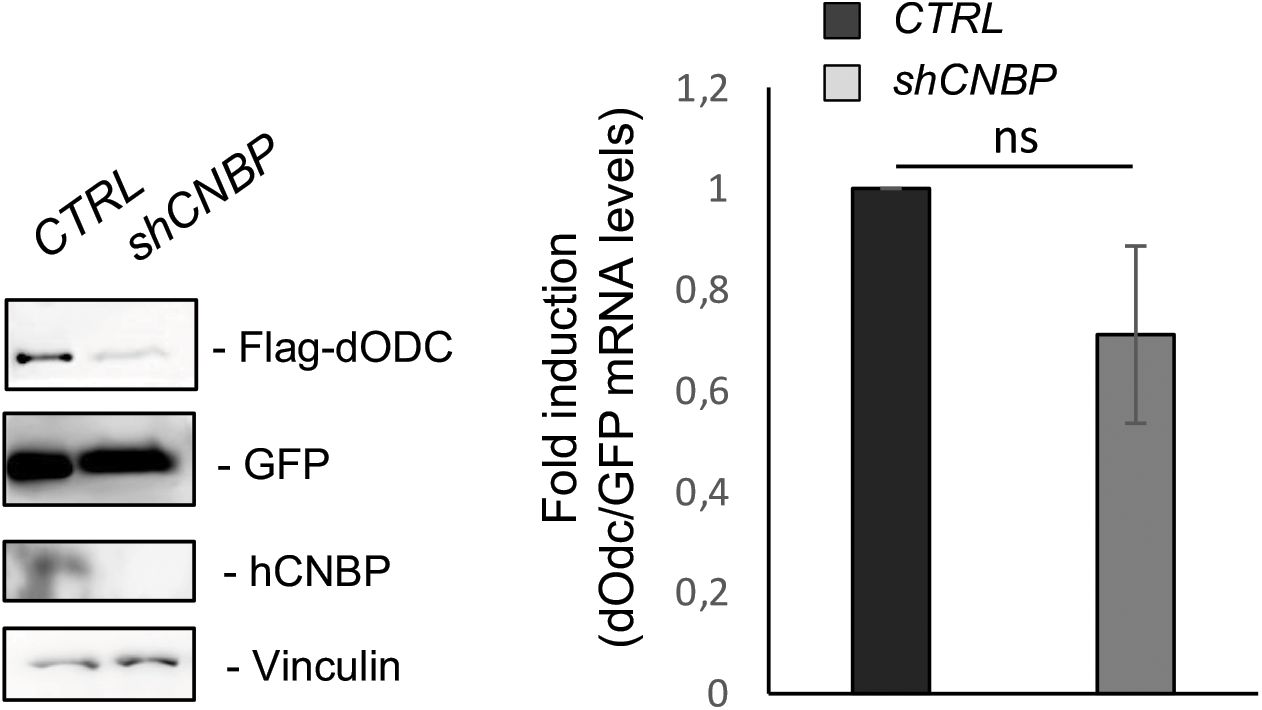
CNBP promotes translation of *dOdc* mRNA. Effect of *hCNBP* depletion on both dODC protein expression and mRNA level of in 293T human cells. On the left the immunoblot shows a direct correlation of hCNBP depletion with a strong reduction of dODC protein levels. Vinculin has been used as loading control. On the right *dOdc1* mRNA levels (qPCR), normalized with the GFP mRNA levels. Error bars represent SEM of three independent experiments; ns: not significative in unpaired t-test.

**Figure 7–figure supplement 1.**
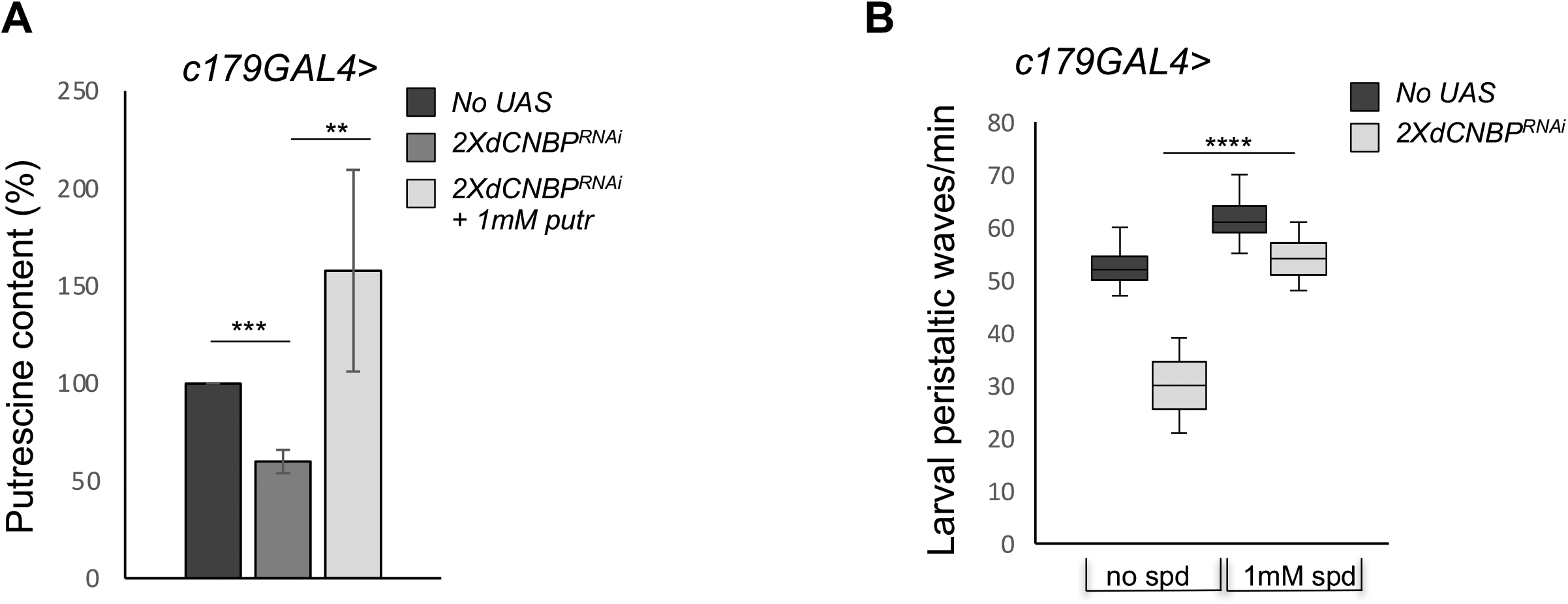
Effects of other polyamines on the CNBP-dependent locomotor phenotype. A. Columns represent the fold difference of putrescine content in dCNBP-depleted larvae grown in absence or presence of 1mM putrescine. *c179GAL4* without putrescine (no UAS; dark grey boxes), *c179GAL4*>2X*UASdCNBP*^*RNAi*^ without 1mM putrescine (dark grey boxes), *c179GAL4*>2X*UASdCNBP*^*RNAi*^ with putrescine (light grey boxes). ** P<0.01, *** P<0.001 with t test. B. Box plot representation of the distribution of peristaltic contraction rates performed by third instar larvae of *c179GAL4* (no UAS; dark grey boxes), *c179GAL4*>2X*UASdCNBP*^*RNAi*^ with or without 1mM spermidine (light grey boxes). The line inside the box indicates the median for each genotype and box boundaries represent the first and third quartiles; whiskers are min and max in the 1.5 interquartile range (**** P<0.0001, ordinary one-way ANOVA post-hoc Tukey’s test). ≥ 10 larvae tested for each genotype in at least two independent experiments. Pool of 10 larvae has been tested for each genotype in two independent experiments. Error bars represent SEM.

**Figure 7–figure supplement 2.**
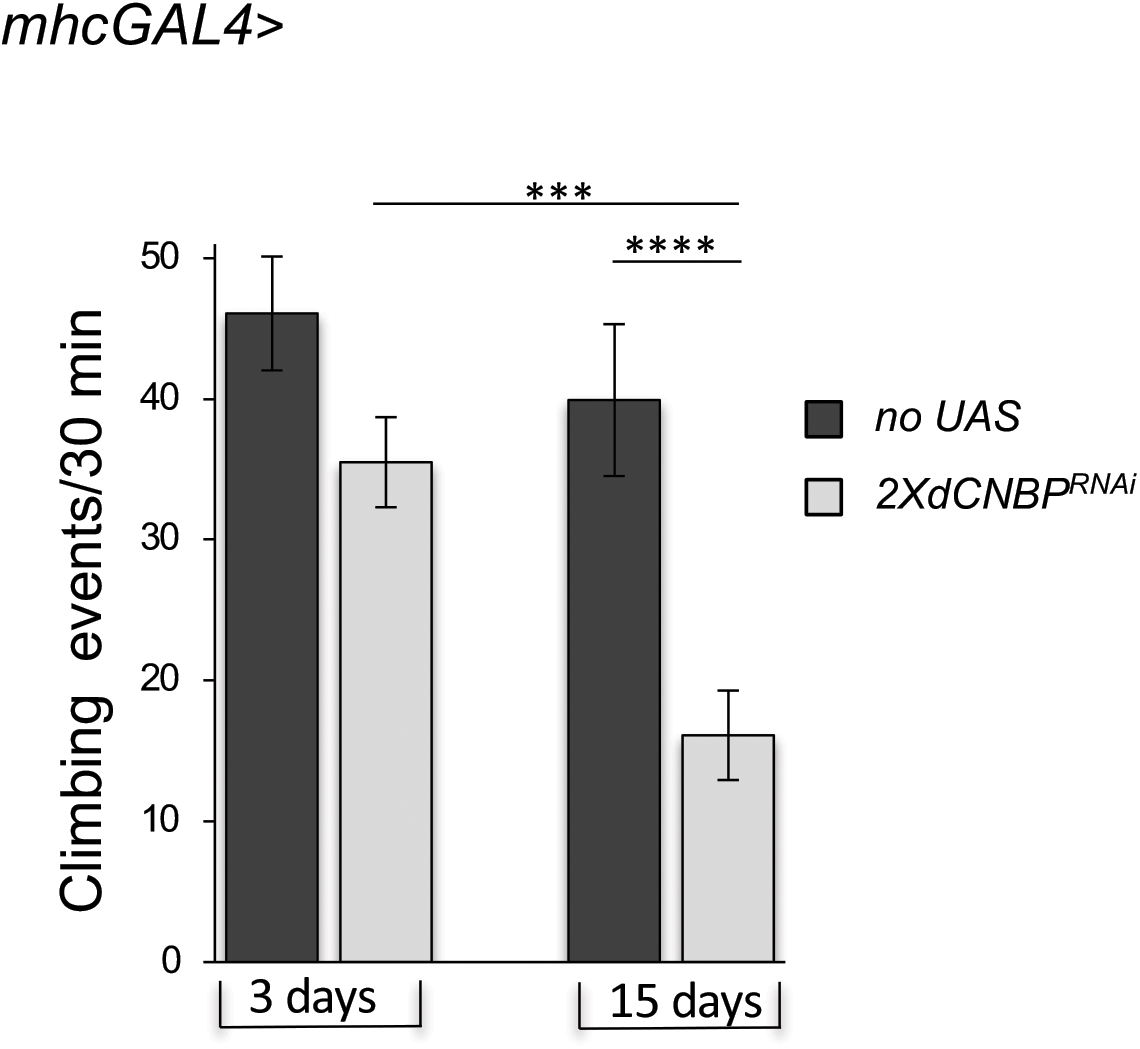
*dCNBP* depleted flies exhibited an age-dependent locomotor dysfunction. *mhcGAL4* or *mhcGAL4 >*2X*UASdCNBP*^*RNAi*^ (29°C) male flies were assayed for negative geotaxis measured by the DAM system at the indicated ages. (Error bars represent SEM; ***p<0.001; ****p<0.0001 Kruskal-Wallis with post-hoc Dunn’s test). ≥ 15 males tested for each genotype in at least two independent experiments.

**Figure 7–figure supplement 3.**
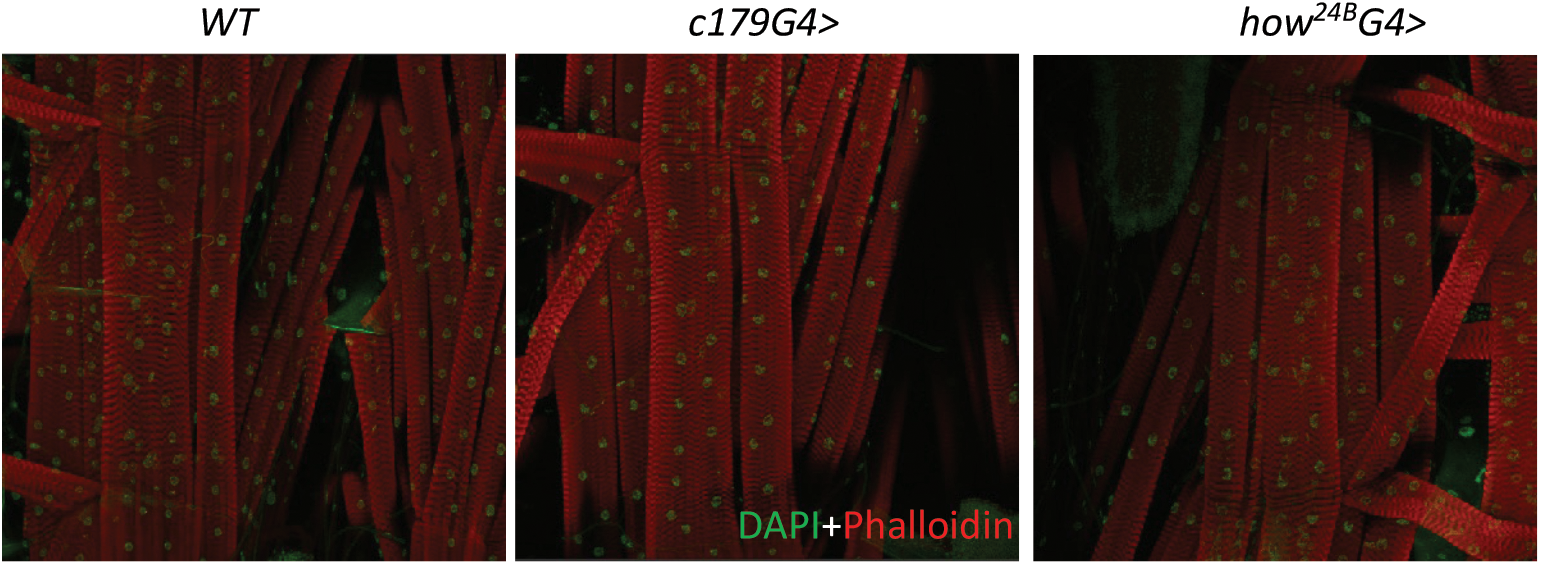
dCNBP depletion does not cause morphological changes of fly muscle tissues. Muscle morphology and sarcomeric organization of the L2/L3 larval body wall muscles. Confocal images of larval muscles (segment A2; muscle 6/7) form *wild type* controls (*WT*) and *2XUASdCNBP*^*RNAi*^ (driven by either *how*^*24B*^*GAL4* or *c179GAL4* as indicated) stained with DAPI (green) and FITC phalloidin (red). ≥ 3 larvae have tested for each genotype in at least two independent experiments.

